# Cortical inhibitory parvalbumin interneurons exhibit metabolic specializations coordinated by PGC-1α that are lost in rodents and humans after traumatic brain injury

**DOI:** 10.1101/2024.06.19.599637

**Authors:** Sadi Quiñones, Jason C Wang, Elliot Kim, Samantha Bottom-Tanzer, Moritz Armbruster, Jéssica K A Macêdo, Tara Hawkinson, Roberto Ribas, Pankaj K. Singh, Lei Wu, Alex R Cantrell, Parveen Kumar, Joshy George, Albert Tai, Michael Whalen, Michael J McConnell, Ramon C Sun, Matthew S Gentry, Chris G Dulla

## Abstract

Parvalbumin-positive interneurons (PV-INs) regulate neuronal and circuit activity, and their dysfunction is observed across neurological conditions, including traumatic brain injury (TBI), epilepsy, Alzheimer’s disease, and schizophrenia. PV-INs are particularly vulnerable to cell loss, potentially due to their increased metabolic demands arising from their uniquely high level of electrical activity, which render them susceptible to metabolic pressure. Here, we use single-nucleus RNA-sequencing (snRNAseq) data from a rodent model of TBI, as well as human TBI data, and demonstrate PV-INs have unique metabolic specializations that are lost after injury and can be rescued by *in vivo* treatment with the glycolytic inhibitor, 2-deoxyglucose. We generated a novel PV-IN transcriptional identity module comprised primarily of genes encoding specialized ion channels, metabolic enzymes, and synaptic machinery, that identifies heterogenous subsets of injury-associated PV-INs with loss of PV-IN transcriptional identity. We show that changes in metabolic specialization are coupled to changes in transcriptional identity in PV-INs and implicate the PV-IN-enriched transcriptional co-activator, *Ppargc1a*, as a key driver of PV-IN transcriptional metabolic dysfunction. We also identify a family of long non-coding RNAs enriched in this subset of transcriptionally dysfunctional PV-INs that negatively correlates with PV-IN metabolic specialization. Lastly, we utilize these tools to interrogate a published human TBI snRNAseq data set and find nearly identical changes, underscoring the importance of PV-IN metabolic dysfunction in the pathology of TBI.

## Introduction

Loss or dysfunction of parvalbumin-positive interneurons (PV-INs) is implicated in the pathophysiology of numerous neurological conditions, including traumatic brain injury (TBI), epilepsy, Alzheimer’s disease (AD), and schizophrenia^1–7^. PV-INs are fast-spiking GABAergic interneurons that control network activity, regulate oscillatory behavior, and ultimately, play an important role in shaping behavior and cognition^3,8–10^. Due to their large excitatory drive and high rate of action potential generation, PV-INs are thought to have heightened energy demands that render them particularly vulnerable to metabolic pressures^3^. While it is known that these neurons express specialized machinery to handle the added stress that is coupled to their function, such as selective expression of the perineural net^11^ (PNN; a specialized form of the extracellular matrix) and *Cox6a2*^12^ (an enzyme involved in oxidative phosphorylation), how these specialized systems fail in neurological disease and render PV-INs uniquely vulnerable is unknown.

In TBI, PV-INs are known to be disproportionately lost and contribute to synaptic, circuit, and behavioral dysfunction^2,4,13–17^. Mounting evidence suggests that loss of PV-INs and inhibitory drive precedes and drives overall network dysfunction, leading to later development of chronic neurological disability such as posttraumatic epilepsy, dementia, and cognitive losses^15–19^. Interestingly, strategies to improve PV-IN function and survival after TBI reduce many injury-associated pathologies in animal models, underscoring the therapeutic potential of PV-IN preservation^13,20–22^. We previously showed that 2-deoxyglucose (2DG), a non-metabolizable glucose analogue and small molecule mimic of the ketogenic diet, administered for 7 days after controlled cortical impact (CCI), a rodent model of contusional TBI, attenuates injury-induced loss of PV-INs, inhibitory dysfunction, and cortical circuit hyperexcitability^13^. Moreover, using genetically labelled PV-INs, we determined that CCI causes both frank PV-IN cell loss, as well as loss of PV protein in surviving, but presumably dysfunctional, PV-INs within the peri-lesional cortex, that was attenuated by in vivo 2DG treatment. Together, this suggests metabolic manipulation of PV-INs may improve both survival and function of PV-INs after TBI, since loss of parvalbumin (PV), a canonical marker and identity molecule of PV-INs, disrupts their function^3,9^.

Here, we use single-nucleus RNA-sequencing (snRNAseq) of peri-lesional cortex 3 days after CCI or sham (uninjured) surgery in male mice to identify multiple metabolic transcriptional modules that are uniquely enriched in PV-INs, are lost after CCI, and are restored by in vivo treatment with 2DG. We identify multiple transcription factors (TFs), including *Esrrg*, *Ncoa2*, *Pparg*, and *Hnf4a*, as coordinators of GABAergic interneuron (GABA-IN) metabolic gene specialization, and validate *Ppargc1a*, a PV-IN-enriched transcriptional co-activator, as a PV-IN-specific regulator of injury-induced metabolic dysfunction. By leveraging injury-induced changes in *Pvalb* (gene encoding PV) expression, we generate a PV-IN transcriptional identity module of ≈ 500 genes whose expression is correlated with *Pvalb* in cortical PV-INs and show that CCI causes bifurcation of PV-INs into Pvalb-ID high and low cells. We identify a set of long non-coding RNAs (lncRNAs), including the inflammation-associated gene, *Miat*, enriched in Pvalb-ID low PV-INs that negatively correlate with GABA-IN-specific metabolic specialization, *Ppargc1a* expression, and PV-IN transcriptional identity. Finally, we re-analyze a published human TBI snRNAseq dataset^23^ and find nearly identical changes to PV-IN biology, including TBI-induced loss of PV-IN transcriptional identity, loss of GABA-IN-specific metabolic specialization, loss of *PPARGC1A*, and increased *MIAT* expression. Together, our findings demonstrate novel transcriptional metabolic specialization in PV-INs, implicate select upstream TFs and PV-IN-specific genes involved in TBI-induced IN dysfunction, and suggest that metabolic intervention may protect PV-IN transcriptional identity, function, and survival in neurological disease.

## Results

### CCI induces early loss of parvalbumin (PV) protein immunoreactivity

Loss of PV-INs occurs after TBI, but when this begins is unknown^13,16,24^. Male C57 mice underwent either moderate CCI or sham surgery, followed by daily IP injections of either 2DG (250 mg/kg) or vehicle saline^13,16^. Three days later animals were sacrificed, brains were prepared for immunohistochemistry, and parvalbumin (PV) protein immunoreactivity was quantified in healthy and injured tissue. Because loss of PV-IN is focal after injury^13^, we quantified PV+ cell density in increments moving away from the site of injury. We found that CCI significantly reduced the density of PV+ cells 3 days after injury, relative to sham, across the entire 500 μm assayed (Fig. 1a-b). This effect was most pronounced near the injury site but occurred throughout the area examined (Fig. 1a-b). The loss of PV immunoreactivity we report here (3 days post injury, dpi) was greater and more spatially broad than in a previous study^13^ examining 2-3 weeks post injury (wpi). While 2DG treatment did not affect PV expression at 3 dpi (Fig. 1a-b), we previously showed treatment with 2DG for 7 days after CCI does increase PV+ density when assayed 2-3 wpi^13^. Together, this suggests that loss of PV immunoreactivity is more extensive at 3dpi compared to 2-3 wpi, and 2DG’s ability to preserve PV-INs chronically might rely on longer time-scale processes such as transcriptional control.

**Figure 1.**
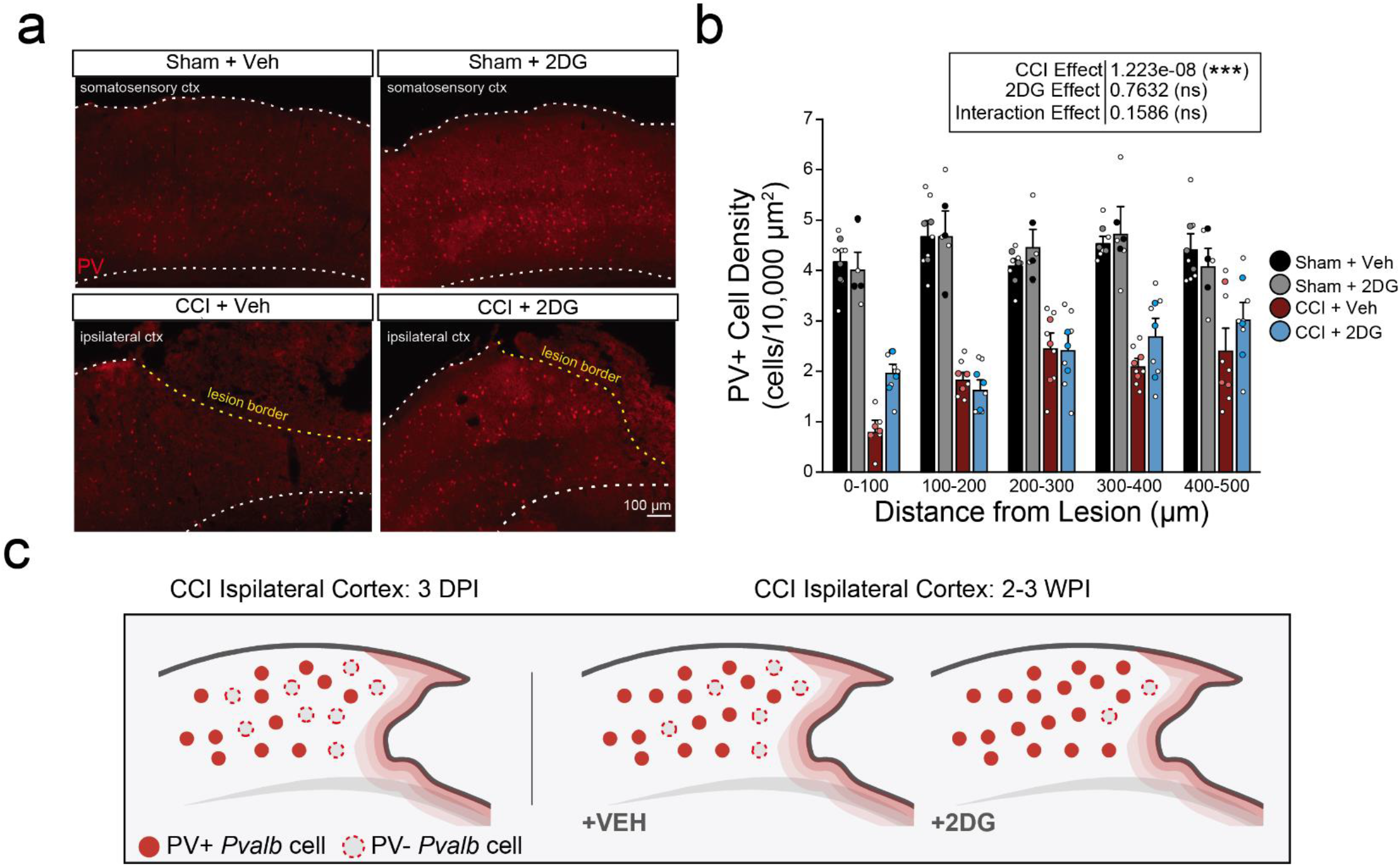
CCI causes significant loss of PV immunoreactivity 3 days post injury. **a**. Representative images of cortical brain sections labeled for parvalbumin (PV, red) immunoreactivity from all experimental conditions (sham + vehicle treatment, sham + 2DG treatment, CCI + vehicle treatment, and CCI + 2DG treatment), 3 days post-injury (dpi). Dotted yellow line represents the lesion border; white dashed line represents the pial surface (top) and white matter border (bottom). Scale bar = 100 µm. **b**. Barplot depicting the density of PV+ cells per 10,000 µm^2^. PV+ cell density is binned in 100 µm bins moving out from the lesion border. Error bars = SEM. *** p = 1.223 e-08, LMM. **c**. Cartoon depiction of PV+ cell density in perilesional cortex at 3 dpi and 2-3 weeks post-injury (wpi) based on this study (3 dpi) and a published study^13^ (3 wpi). Dark red filled circles represent PV+ cells and grey filled dotted circles represent PV immuno-negative *Pvalb* cells.

### Single nucleus profiling of CCI-injured and 2DG-treated cortex at 3 days after injury

To determine how 2DG affects the PV-IN response to injury, and to ask what metabolic differences might render PV-INs uniquely susceptible to neurological insult compared to other neuronal subtypes, we performed snRNAseq on cortical brain tissue from CCI and Sham animals treated with either vehicle saline or 2DG. Three days after CCI, perilesional cortical tissue was collected and, in shams, analogous cortical tissue was collected. Care was taken to exclude underlying white matter, as well as cortical tissue distal to the injury lesion (see Methods; Supp. Fig. 1). Using the 10x Genomics Chromium platform, we generated 12 cDNA libraries across 4 groups: Sham+Vehicle, Sham+2DG, CCI+Vehicle, and CCI+2DG. Three independent replicates of each group were generated, each containing samples pooled from three mice. After quality control (see Methods), we sequenced 92,589 nuclei across 36 mice in 4 groups (Sham+Veh: 21,967 nuclei; Sham+2DG: 21,576 nuclei; CCI+Veh: 23,056 nuclei; CCI+2DG: 25,990 nuclei) (Fig. 2f). We integrated our datasets using CCA^25^ and clustered nuclei based on their integrated gene expression profiles (Fig. 2a). Cluster identities were then annotated based on established cell type-selective gene expression (Fig. 2a,d) and neurons were further sub-clustered (Fig. 2a-b,d,h). Glutamatergic principal neurons (Glut-PN) clusters were annotated based on layer-specific markers and further sub-divided into transcriptionally distinct subpopulations (Fig. 2d,h). GABAergic INs (GABA-INs) were clustered into 2 groups corresponding to their developmental origins in the medial (MGE) or caudal ganglionic eminence (CGE) (Fig. 2b). Within the CGE-INs, two clusters were identified: VIP-INs (*Adarb2*, *Vip*), and LAMP5-INs (*Adarb2*, *Lamp5*) (Fig. 2d). In the MGE-INs, we identified an SST-IN cluster (*Lhx6*, *Sst*) and PV-IN cluster (*Lhx6*, *Pvalb*) (Fig. 2d).

**Figure 2.**
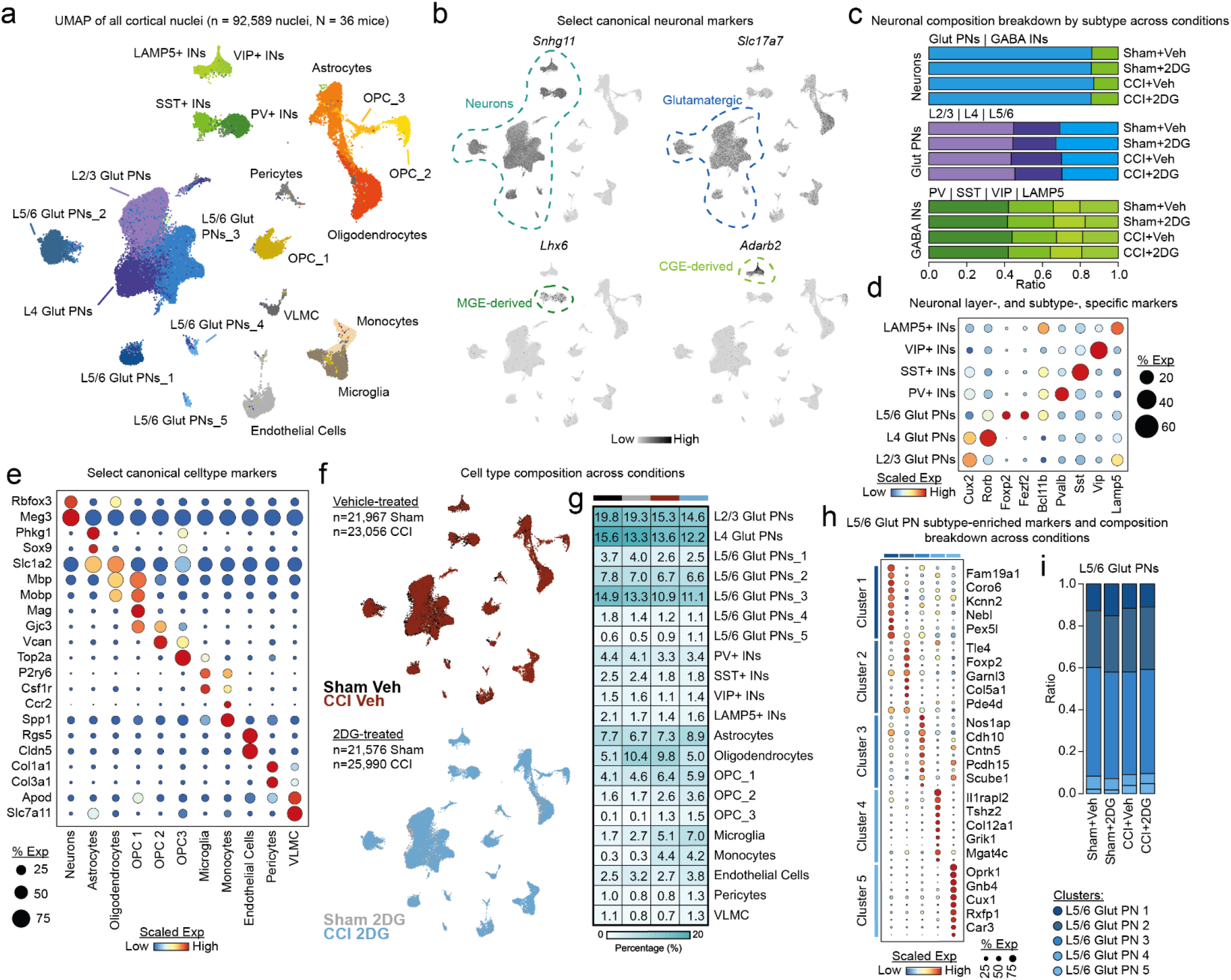
Single nucleus RNA sequencing of CCI and 2DG-treated cortex captures all major cell types. **a.** UMAP depiction of all nuclei captured, colored by cell type cluster identity. **b.** Various UMAPs of nuclei colored by scaled expression of select neuronal marker genes – *Snhg11, Slc17a7, Lhx6, Adarb2*. Each gene is significantly enriched in dotted circled clusters, p < 0.0001 by Wilcoxon Rank Sum with Bonferroni correction. **c.** Stacked barplots illustrating the compositional breakdown of neuronal subtypes by category (neurons, Glut-PNs, and GABA-INs) across experimental conditions. Colors represent neuronal subtypes. **d.** Dotplot depicting scaled expression of neuronal layer-, and subtype-specific marker genes across neurons. **e.** Dotplot depicting scaled expression of select canonical cell type marker genes across all cells. **f.** UMAP depiction of all cells captured in vehicle-treated datasets - Sham+Veh, CCI+Veh (*top*). UMAP depiction of all cells captured in 2DG-treated datasets - Sham+2DG, CCI+2DG (*bottom*). **g.** Heatmap illustrating the percent of total nuclei that each cluster comprises, split by condition. **h.** Dotplot depicting scaled expression of top 10 DEGs in each L5/6 Glut-PN cluster, p < 0.0001 by Wilcoxon Rank Sum with Bonferroni correction. **i.** Stacked barplots illustrating the compositional breakdown of L5/6 Glut-PNs.

In line with published data, our total neuronal population was comprised of approximately 86% Glut-PNs and 14% GABA-INs (Fig. 2c)^8,26,27^. GABA-INs consisted of 66% MGE-INs and 34% CGE-INs and consistent with cortical GABA-IN diversity studies, our PV-IN cluster comprised roughly 42% of the entire GABA-IN population (Fig. 2c)^8,26,27^. Given the injury context, we next evaluated the change in cellular composition across conditions. Overall, we captured relatively similar proportions of each cluster identity across conditions, particularly within neuronal subtypes (Fig. 2c,g,i). Exceptions include CCI-enriched cell types such as microglia, infiltrating monocytes, transition-state oligodendrocyte precursor cells, *Vcan*+ OPC 2, and proliferative oligodendrocyte precursor cells, *Top2a*+ OPC 3, (Fig. 2g)^28,29^. These results show that our snRNAseq approach successfully captured all major cell types of the brain, including CCI-specific cell types and physiologically analogous proportions of neuronal subtypes.

### Cell type-specific injury-, and 2DG-, associated transcriptional pathways

Next, we defined CCI-, and 2DG-, associated changes by identifying differentially expressed genes (DEGs) between Sham+Veh and CCI+Veh cell types, as well as between CCI+Veh and CCI+2DG cell types (Methods; Fig 3a). There was a modest, but significant correlation between the number of nuclei per group and the number of DEGs identified (Supp. Fig. 2a). This was eliminated when neuronal populations were merged into higher order groupings (Supp. Fig. 2b), so subsequent analysis used larger order groupings (Glut-PNs grouped by layer, GABA-INs grouped by developmental origin). We then performed gene set enrichment analysis (GSEA) on DEGs (inclusion criteria: min.pct > 0.1, p < 0.1) to identify pathways that were significantly up-, and down-, regulated (Fig. 3a)^30^. By then ranking the GSEA pathways based on statistical significance, we identified the most relevant changes associated with CCI and 2DG treatment. We found that many of the top affected pathways were conserved across all neuronal subtypes and consistent with previous snRNAseq studies (Fig. 3b)^29,31–35^. Oxidative Phosphorylation and Mitochondrial Dysfunction were the most significantly affected pathways across all cells, especially neurons, after CCI, consistent with metabolic dysfunction as a primary pathology of TBI. When we compare the top 25 CCI-associated GSEA terms to the top 25 2DG-associated GSEA terms, we find that many of the top pathways are conserved but in the opposite z-score direction (Fig. 3b-c), consistent with 2DG reversing CCI-induced transcriptional changes. eIF2/eIF4 signaling and mTOR pathways were enriched in 2DG-associated GSEA after CCI, consistent with known 2DG effects following stroke^36^.

**Figure 3.**
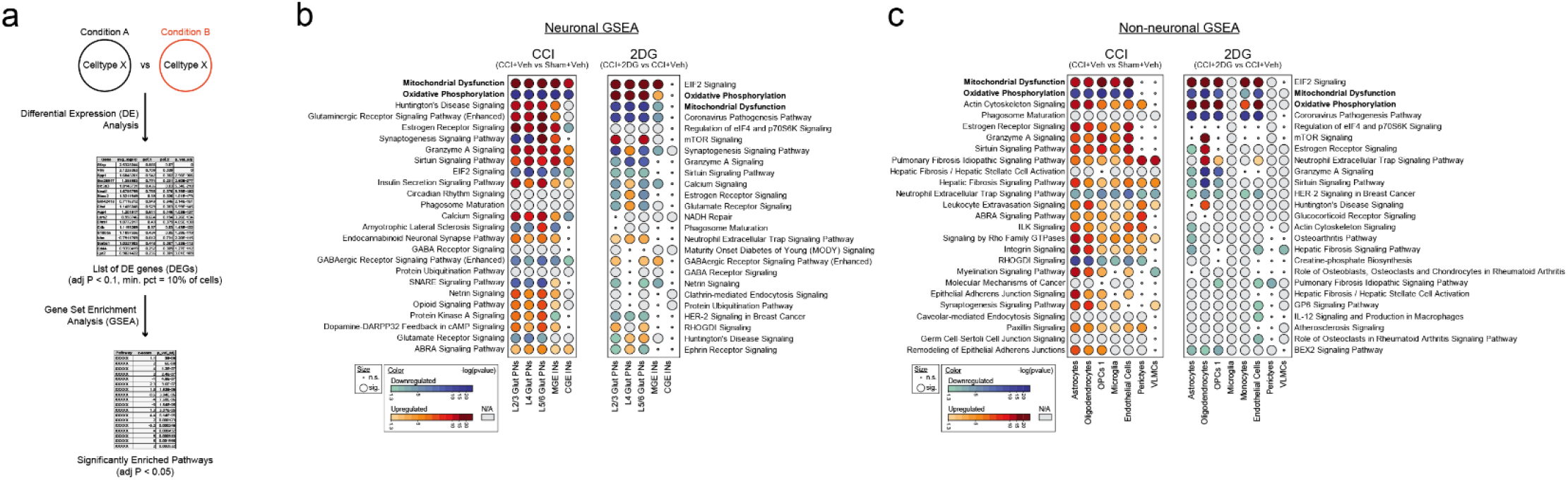
CCI-, and 2DG-, associated changes to the single cell transcriptome. **a**. Workflow schematic depicting scheme for pathway analysis. Cell types were compared across conditions and DE analysis was performed. DEGs were then processed using GSEA in Ingenuity Pathway Analysis (IPA) to identify significantly enriched pathways in each cell type. **b**. Heatmap illustrating the top 25 most significantly enriched pathways affected by CCI (left) and 2DG (right) in neuronal populations identified by GSEA in each sub type. Red = upregulated; Blue = downregulated; significant changes are shown by large circles. **c**. Same as (**b.**), but for non-neuronal populations.

We next compared these results to gas chromatography mass spectrometry (GC-MS) metabolomic analysis of perilesional cortical tissue 3 dpi. We resolved the relative concentrations of 46 total metabolites (glycolytic intermediates, TCA metabolites, and amino acids; Supp. Fig. 3); 16 of which were significantly affected by injury (p < 0.05, two-way ANOVA with FDR correction). Surprisingly, despite the large effect of 2DG on metabolic gene expression, it did not significantly alter global metabolomic changes caused by TBI (Supp. Fig. 3). Overall, this supports the hypothesis that metabolic dysfunction is a primary aspect of neuronal and non-neuronal dysfunction in the days following CCI. It also suggests that, although 2DG acts to promote increased oxidative phosphorylation and mitochondrial gene expression, it does not change bulk metabolomic state.

### GABA-INs have enriched metabolic gene expression that is compromised by CCI and restored by 2DG

Bulk metabolomics did not reveal significant global changes caused by 2DG, and CCI-associated GSEA showed similar metabolic disruption across all neuronal subtypes; yet PV-Ins are distinctly susceptible to loss after CCI and preserved by 2DG. Thus, we next asked whether underlying differences in neuronal subtype-specific metabolic gene expression might underlie differences in response to injury and 2DG treatment. We first characterized typical cell type-specific metabolic gene expression. We assessed 1467 metabolism-related genes from the KEGG database^37^ encompassing canonical energy-producing pathways (e.g. glycolysis, TCA), as well as other types of metabolism (e.g. RNA and amino acid metabolism). Genes not captured in our data were removed, leaving 1198 metabolic genes that we used to perform DE analysis between different cell types within the Sham+Veh dataset (Fig. 4a). We grouped our cells by their higher order groupings (neurons, glia, and vasculature) and monocytes. *Vcan*+ OPC 2, and *Top2a*+ OPC 3 groups were excluded given their rare (<1%) appearance in Sham+Veh datasets. This revealed a significant enrichment of specific genes in each cell type, with notable difference between neurons and other cells (Fig. 4a). When we examined the top 100 neuron-enriched metabolic genes, we observed laminar-, and subtype-, specific distribution of metabolic genes (Supp. Fig. 4).

**Figure 4.**
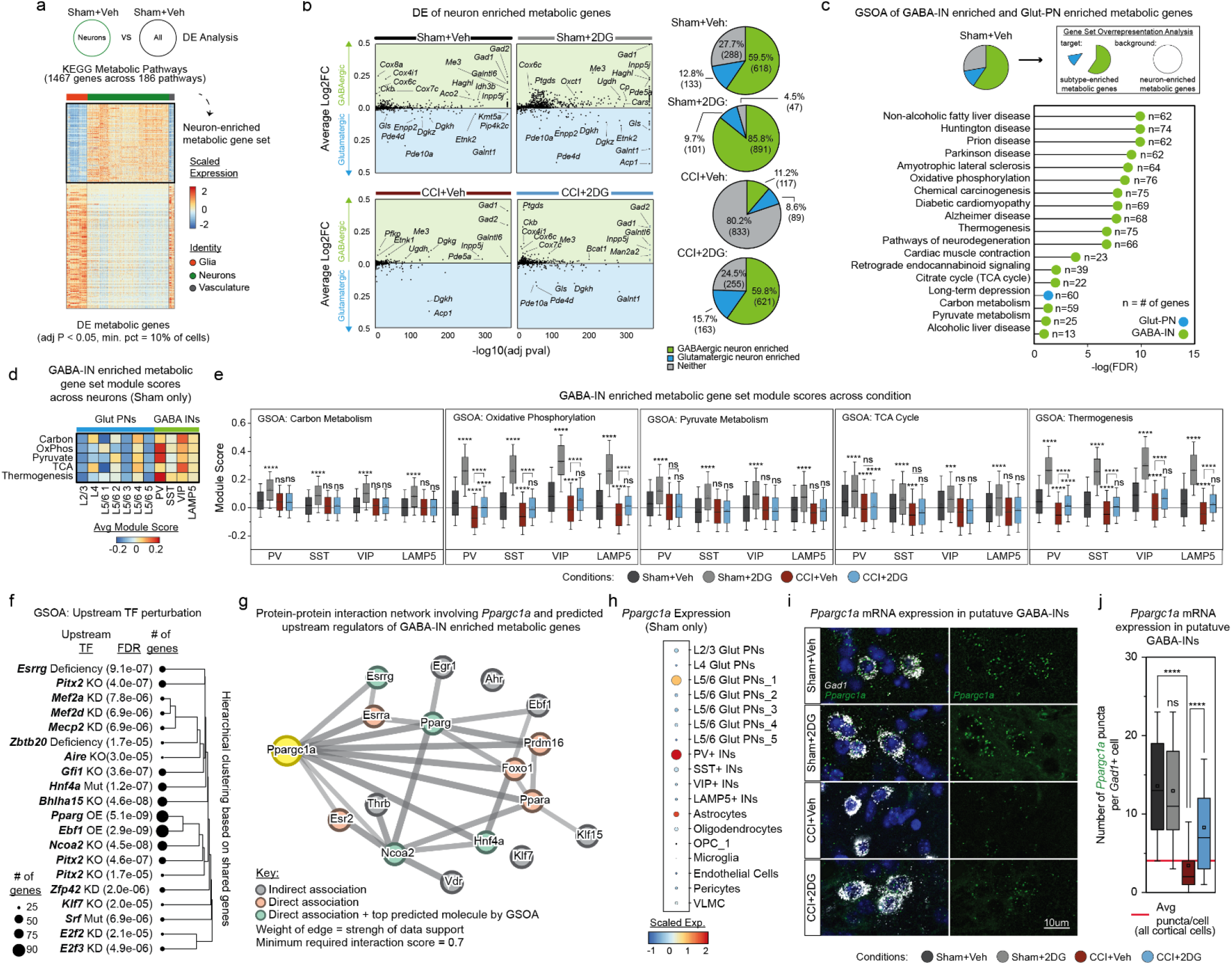
Metabolic gene expression is enriched in GABA-INs, is lost after TBI, is restored by 2DG treatment, and correlates with PGC1-α expression. **a.** Schematic depicting workflow used to identify neuron-enriched metabolic genes. Heatmap shows scaled expression of metabolic genes (rows) across all cells (columns) from the Sham+Veh dataset. Glia group includes astrocytes, oligodendrocytes, microglia, and OPCs. Neuron group includes GABA-INs and Glut-PNs. Vasculature group includes endothelial cells, pericytes, and VLMC. **b.** Scatter plots and pie chart quantification showing the relative enrichment of each neuron-enriched metabolic gene determined in (**a.**) in either GABA-INs (green zone) or Glut-PNs (blue zone). Quantification of GABA-IN vs Glut-PN enrichment determined by DE analysis on neuron-enriched metabolic genes comparing all GABA-INs to all Glut-PNs (Wilcoxon Rank Sum test with Bonferroni correction, adj P < 0.05). **c.** Lollipop graph displaying the exhaustive list of significantly overrepresented pathways corresponding to enriched metabolic genes in GABA-INs (green) and Glut-PNs (blue) from (**c.**); FDR < 0.05. n = number of genes in dataset. **d.** Heatmap showing the module score for each GSOA term identified in (**c.**) across each neuronal subtype in Sham+Veh. **e.** Boxplots of the module scores for each GSOA term across each GABA-IN subtype in all experimental conditions. **** p < 0.0001 by Wilcoxon Rank Sum test with Bonferroni correction. **f.** Tree plot illustration of the top predicted upstream TFs based on GABA-IN-enriched metabolic genes altered by TBI and restored by 2DG. Size of circle corresponds to number of shared genes. **g.** Protein-protein interaction network of TF identified in (**f.**). Nodes consist of *Ppargc1a* and 16 genes identified as predicted upstream regulators of GABA-IN metabolic enrichment in (**f**.). **h.** Dotplot of the average *Ppargc1a* mRNA expression in each cell type in Sham+Veh from snRNAseq. **i.** Representative *Ppargc1a* smFISH images from all experimental conditions. CCI images were taken within the first 500 µm of perilesional cortex in layer 5/6. Sham images were taken in layer 5/6 somatosensory cortex. *Gad1* (white) and *Ppargc1a* (green). Scale bar = 10µm. **j.** Boxplot of number of *Ppargc1a* mRNA puncta per *Gad1*+ cell across experimental conditions; inner squares represent mean values; inner black lines represent median values. Lower and upper box limits represent 25th and 75th percentile, respectively. Error bars represent 10th and 90th percentile. Red line marks the average number of Ppargc1a puncta per cell in all cells. **** p <0.0001 by Wilcoxon Rank Sum test with Bonferroni correction.

We next examined if neuron-enriched metabolic genes were preferentially expressed in excitatory (Glut-PNs) or inhibitory (GABA-INs) neurons by performing DE analysis on the same set of metabolic genes. We found that 59.5% of neuronally enriched metabolic genes were enriched in GABA-INs, compared to 12.8% of genes enriched in Glut-PNs, while 27.7% of genes were not enriched in either group (Fig. 4b). We then used Gene Ontology Overrepresentation Analysis (GSOA)^38^ of GABA-IN-enriched metabolic genes to identify 15 significantly enriched pathways. 5/15 pathways belonged to energy-producing metabolic processes (TCA cycle, oxidative phosphorylation, thermogenesis, carbon metabolism, and pyruvate metabolism), while many neurological disease-associated pathways (AD, HD, PD, and ALS) were also identified (Fig. 4c). GSOA of Glut-PN-enriched metabolic genes identified only 1 significantly enriched pathway, long term depression (Fig. 4c). As a control to test whether metabolic enrichment in GABA-INs was biological, rather than a sampling artifact, we tested whether the observed metabolic enrichment signature was an outlier compared to random gene sampling. We generated an enrichment signature for 30 individual, randomly generated gene sets comprised of 1000 non-repeating genes. We then used a Grubbs test to assess whether the GABA-IN-enriched metabolic gene set was a statistical outlier compared to randomly sampled genes (Supp. Fig. 5a-b). Grubbs test confirmed that the metabolic gene set was an outlier, supporting our finding that GABA-INs are metabolically enriched as a biological effect, rather than sampling artifact (Supp Fig 5a-b).

**Figure 5.**
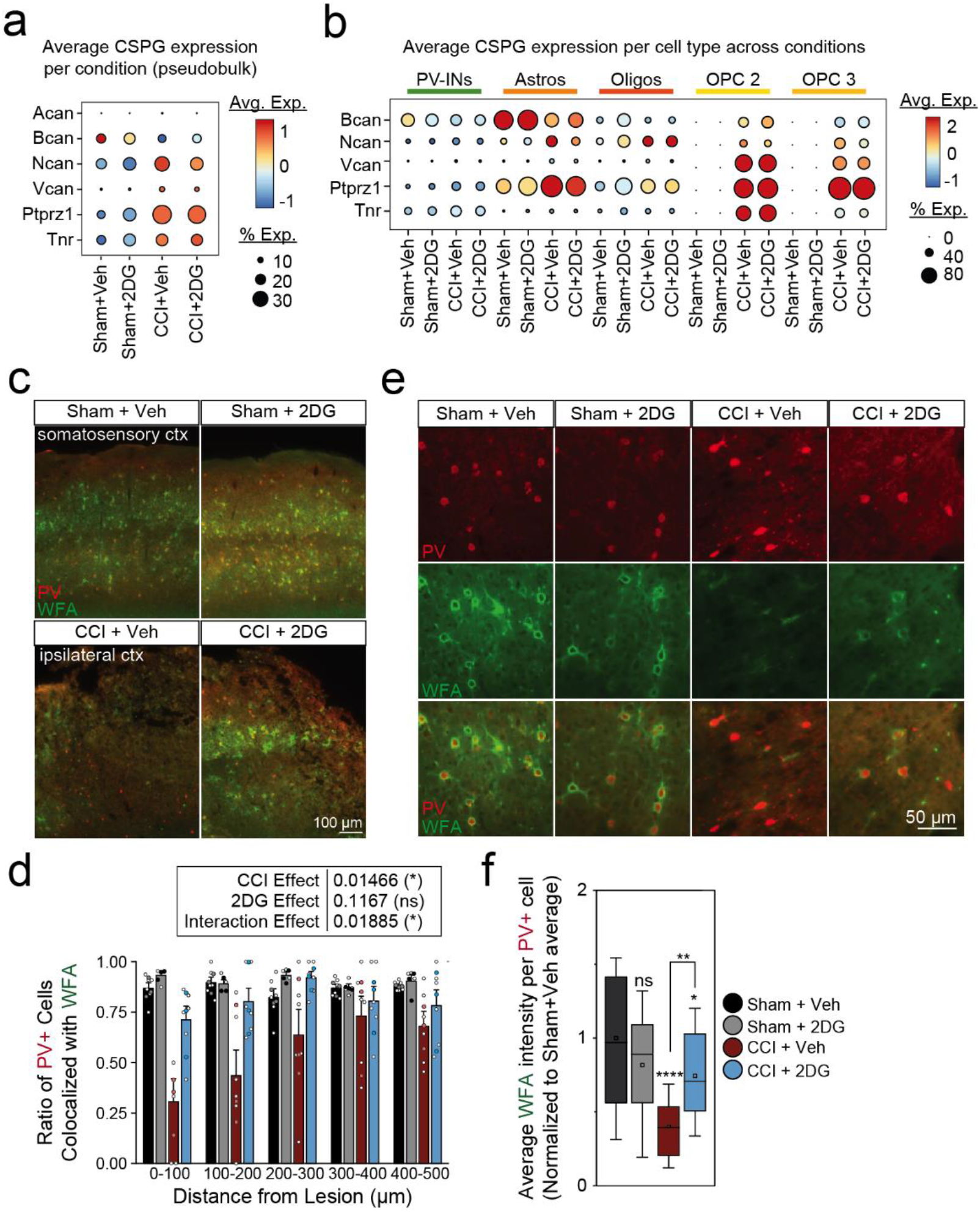
2DG rescues CCI-induced disruption of PNN, a specialized ECM, surrounding PV-INs. **a.** Dotplot of average expression of various PNN-related chondroitin sulfate proteoglycans (CSPGs) per condition. Each condition is a pseudobulk representation. **b.** Dotplot of average expression of CSPGs from (**a.**) per cell type relevant to PNN CSPG production. **c.** Representative PV immunolabeled (red) and WFA labeled (green) images from all experimental conditions, 3 dpi. Scale bar = 100 µm. **d.** Barplot depicting ratio of ‘PNN+ PV+’ cells to ‘PNN-PV+’ cells per 100 µm from injury border. Error bars = SEM. * p < 0.05, LMM **e.** Higher magnification images of PV immunolabel and WFA stain of PNN. Scale bar = 100 µm. **f.** Boxplot of the mean intensity of WFA per PV+ cell normalized to the average Sham+Veh value. Inner squares represent mean values, inner black lines represent median values. Lower and upper box limits represent 25th and 75th percentile, respectively. Error bars represent 10th and 90th percentile. * p < 0.05, ** p < 0.01, **** p < 0.0001 by two-way ANOVA with Bonferroni correction. Statistical comparisons are made relative to Sham+Veh, unless otherwise denoted by brackets.

We next tested whether CCI or 2DG treatment altered metabolic gene enrichment observed in Sham+Veh GABA-INs. We performed DE analysis on the same neuron-enriched metabolic gene set across all experimental conditions and found a substantial decrease in GABA-IN-enriched metabolic genes following CCI (59.5% of genes were enriched in GABA-INs in Sham+Veh versus 11.2% in CCI+Veh; Fig. 4b). Conversely, 2DG treatment increased the number of enriched metabolic genes in both CCI and Sham conditions (Fig. 4b). To address whether the increase in metabolic enrichment of GABA-IN by 2DG represents a restoration of the Sham+Veh gene enrichment signature, rather than a different set of metabolic genes, we plotted the average enrichment of each gene from Sham+Veh against the enrichment of those genes across the other conditions. A linear fit was then applied, the slope of the fit was used to determine the average level of metabolic enrichment relative to Sham+Veh, and a Pearson correlation was calculated to test the strength of that relationship. The Sham+Veh ∼ CCI+2DG correlation analysis revealed a surprisingly strong linear fit of the data with a slope near 1 (Pearson r = .90, R2 = .81), supporting a restoration of the original GABA-IN metabolic gene enrichment to Sham+Veh levels and not a new pattern of metabolic gene expression (Supp. Fig. 6a-b).

We next generated gene set modules^39^ using the GABA-IN-enriched metabolic genes that pertain to each GSOA term identified (Methods). This allowed us to interrogate specific GABA-IN-relevant metabolic pathways in individual neuronal subtypes and across conditions. This validated that each GSOA module score was significantly enriched in GABA-INs relative to Glut-PNs in the Sham+Veh condition, particularly enhanced in PV-, and to a smaller extent VIP-, INs (Fig 4d). Given these findings, we restricted the remaining analysis to GABA-INs and generated GSOA-term module scores for GABA-INs across conditions. Consistent with GSEA, GSOA showed that injury decreased OxPhos, TCA cycle, thermogenesis, and pyruvate metabolism in PV-INs (Fig 4e) while 2DG had the greatest attenuative effect on OxPhos and thermogenesis (Fig 4e). Together, these findings demonstrate a broad enrichment of 100s of metabolic genes in GABA-INs that is lost after CCI but restored by *in vivo* treatment with 2DG.

### PGC-1**α** is a PV-IN enriched master regulator of metabolism implicated in TBI-induced changes in metabolism

Based on the coordinated shift in GABA-IN metabolic gene enrichment observed after CCI and 2DG treatment, we used GSOA to identify potential upstream regulators of GABA-IN-enriched metabolic genes. We generated a list of ≈ 500 GABA-IN-enriched metabolic genes that were diminished by CCI and enhanced by 2DG and performed GSOA using a reference database of transcription factor (TF) perturbation experiments. This revealed many TFs whose downstream targets significantly overlapped with CCI-, and 2DG-, associated GABA-IN-enriched metabolic genes, including *Prdm16*, *Ebf1*, and *Ncoa2* (Fig 4f). Interestingly, many of these TFs are known to have convergent downstream mechanisms and are ubiquitously expressed across most, if not all, neurons^40,41^. Thus, to identify sub-networks of interacting molecules, based on these predicted upstream TFs, that might confer PV-INs unique metabolic profile, we ran STRING^42^ protein-protein interaction network analysis on the predicted TFs (FDR < 0.05, normalized enrichment scores (NES) < 0.01) (Methods). Clustering of the TFs based on connectivity revealed a cluster of TFs with enrichment for oxidative metabolism, nuclear receptors, and mitochondrial function (Fig. 4g). When we computationally expanded the network of nodes, we identified *Ppargc1a*, a PV-IN-enriched master regulator of metabolism, as the nearest neighbor, highlighting a potential PV-IN-specific mechanism of injury-induced transcriptional dysfunction.

PGC-1α, encoded by the gene *Ppargc1a*, is a master transcriptional coactivator that coordinates metabolic processes including mitochondrial biogenesis, oxidative metabolism, and glucose metabolism^43–46^. Deletion of PGC-1α in PV-INs causes asynchronous GABA release and downregulates key functional PV-IN-specific genes such as *Pvalb* and *Syt2*^45–47^. PGC-1α expression in the brain suggests that it is preferentially enriched in highly metabolically active cell types, including glia, fast-spiking INs such as PV-INs, and large projection neurons^47^. We quantified the average expression of *Ppargc1a* across all cell types in the healthy brain in snRNAseq data and found it was restricted to PV-INs, astrocytes, *Top2a*+ OPC 3, and L5/6 Glut-PN 1 with the highest expression in PV-INs (p < 0.0001 by Wilcoxon Rank Sum with Bonferroni correction) (Fig 4h). To validate GABA-IN enrichment of *Ppargc1a* expression, we prepared cortical sections for single molecule fluorescent in situ hybridization (smFISH). We applied probes against *Sox9*, a marker of astrocytes^48^, *Gad1*, a marker of GABA-INs, and *Ppargc1a*. *Gad1*, a general marker of GABA-INs, was used instead of *Pvalb*, which specifically labels PV-INs, since *Pvalb*, but not *Gad1*, showed significant downregulation after CCI (data not shown). We found that *Gad1*+ putative GABA-INs expressed ≈ 4 times as many *Ppargc1a* mRNA puncta than *Sox9*+ putative astrocytes and *Gad1*-/*Sox9*-cells (Supp. Fig. 7b,c). Taken together, this data supports the evidence that *Ppargc1a* expression is preferentially expressed in PV-INs, and its expression is enriched relative to other *Ppargc1a*+ cell types.

Next, we quantified *Ppargc1a* expression across conditions and found a 75% decrease in the median number of *Ppargc1a* puncta per putative GABA-IN in CCI+Veh compared to Sham+Veh (Sham+Veh median: 13.57, CCI+Veh median: 3.43; p < 0.0001 by Wilcoxon rank sum with Bonferroni correction) (Fig. 4i-j). After CCI, ∼75% of putative GABA-INs have levels of *Ppargc1a* expression below background, consistent with complete loss of expression (Fig. 4j). 2DG treatment after CCI resulted in >2-fold change (FC) (p < 0.001 by Wilcoxon rank sum with Bonferroni correction) in median expression of *Ppargc1a,* with no effect in sham injured animals (−1.05 FC, p = 0.78 by Wilcoxon rank sum with Bonferroni correction) (Fig. 4i-j). These results show *Ppargc1a* is enriched in healthy PV-INs, lost after CCI, and restored by in vivo treatment with 2DG, implicating it as a central regulator of PV-IN-specific metabolic gene expression in the injured brain.

### Oxidative metabolism relates metabolic function to the PNN, a specialized extracellular matrix

Beyond their specialized metabolic capacity, fast-spiking PV-INs are surrounded by a specialized extracellular matrix (ECM), known as the perineural net (PNN)^24,49–51^, which shapes their excitability and synaptic input, and is disrupted in acute and chronic TBI^11,24,49^. PNN influence PV-IN metabolism and, in a reciprocal manner, PV-IN metabolism influences PNN integrity^12^. Because 2DG restores PV-IN metabolic gene expression, we suspected it may also preserve PNN integrity after TBI. Many genes across multiple cell types regulate PNN generation and degradation. Some are PNN-specific, while others are also involved in other ECM structures^49^. Using snRNAseq, we focused on chondroitin sulfate proteoglycans (CSPGs), the main structural components of the PNN (Fig. 5a-b). We found general dysregulation of CSPGs by CCI and 2DG, in a cell type-dependent manner (Fig. 5a-b). Astrocytes, oligodendrocytes, and OPCs showed the most dramatic changes in CSPG expression, particularly in *Ncan* and *Vcan* expression (Fig. 5b) which are key components of the glial scar and OPC responses to injury^28,52^. We focused on brevican (*Bcan*), given its specificity to PNN, and found CCI significantly decreased *Bcan* expression in both PV-INs and astrocytes, the two main cell types responsible for its secretion (Fig. 5a-b)^53,54^. 2DG-treatment modestly increased *Bcan* expression after CCI (Fig. 5a-b).

Next, we performed immunohistochemical validation of changes in PNN using an antibody against PV and *Wisteria floribunda* Agglutinin (WFA), a lectin that labels PNNs. First, we asked whether CCI and 2DG influenced the ratio of PV-INs that were co-localized with PNN. We found that, on average, 87 ± 7.0% (average± standard deviation) of PV-INs were co-localized with PNN in Sham+Veh cortex (Fig. 5c-d). CCI caused a drastic reduction in PNN+ PV-IN ratio relative to Sham+Veh (CCI effect, p < 0.05 by LMM), that was significantly attenuated by 2DG treatment (interaction effect, p < 0.05 by LMM) (Fig. 5c-d). Further, when we quantified the mean intensity of each PNN via WFA labeling, we found CCI caused a ∼50% reduction in mean intensity per PNN relative to Sham+Veh (p < 0.0001 by two-way ANOVA with Bonferroni correction) (Fig. 5e-f) that was largely restored by 2DG treatment (p < 0.01 by two-way ANOVA with Bonferroni correction) (Fig. 5e-f). Because PNN degradation is permissive to changes in synaptic input^49,50^, and loss of PNN during injury has been shown to lead to PV-IN oxidative damage and excitotoxic death^24^, this suggests a potential neuroprotective benefit of 2DG treatment by attenuation of pathological synaptic remodeling surrounding PV-INs.

### Heterogenous PV-IN populations after injury demonstrate relationship between PV-IN identity and oxidative metabolism

CCI causes early loss of PV protein and cellular dysfunction in a subset of PV-INs, particularly those closest to site of injury. With the rationale that these early dysfunctional PV-INs could be targeted for therapeutic intervention, we wanted to characterize the molecular heterogeneity among PV-INs in CCI+Veh. We examined 3,498 PV-INs across all conditions (Fig. 6a) and generated a “Pvalb-Identity” (Pvalb-ID) module comprised of 557 genes whose expression positively correlated with *Pvalb* in PV-INs (Pearson’s r > +0.5) (Fig. 6b). The top Gene Ontology Molecular Function (GO:MF) terms for genes in the Pvalb-ID module were ‘oxidoreduction-driven active transmembrane transporter activity’ and ‘RNA binding,’ which include relevant genes such as *Ppargc1a*, *Tfam*, *Cox* family genes, and *Ndufa* family genes (Fig. 6b)^55–57^. Additionally, this module includes genes critical to PV-IN function (*Kcna1, Kcnc1, Kcnc3, Syt2*, and others). On average, we found that Pvalb-ID module scores were decreased by CCI and partially restored by 2DG in PV-INs (Fig. 6). However, in line with PV-IN heterogeneity, CCI+Veh PV-INs displayed a bimodal distribution of Pvalb-ID module scores (Fig. 6c-d) with one population centered around a Pvalb-ID score of ≈ 0.05 (enriched) and another centered around a score of ≈ −0.04 (de-enriched) (Fig. 6c-d). Based on this enrichment and de-enrichment of identity scores, we split CCI+Veh PV-INs into two groups, Pvalb-ID high (module score > 0.005) and Pvalb-ID low (module score < 0.005). *Pvalb* mRNA expression alone was insufficient to stratify our PV-INs (Supp. Fig. 8)^1,13^. DE analysis comparing Pvalb-ID high and low PV-INs showed significant enrichment of several inflammatory-related long non-coding RNAs (lncRNAs) in Pvalb-ID low PV-INs, including *Miat, Mirg, and Gm26917* (p < 0.0001 by Wilcoxon rank sum with Bonferroni correction) (Fig. 6e).

**Figure 6.**
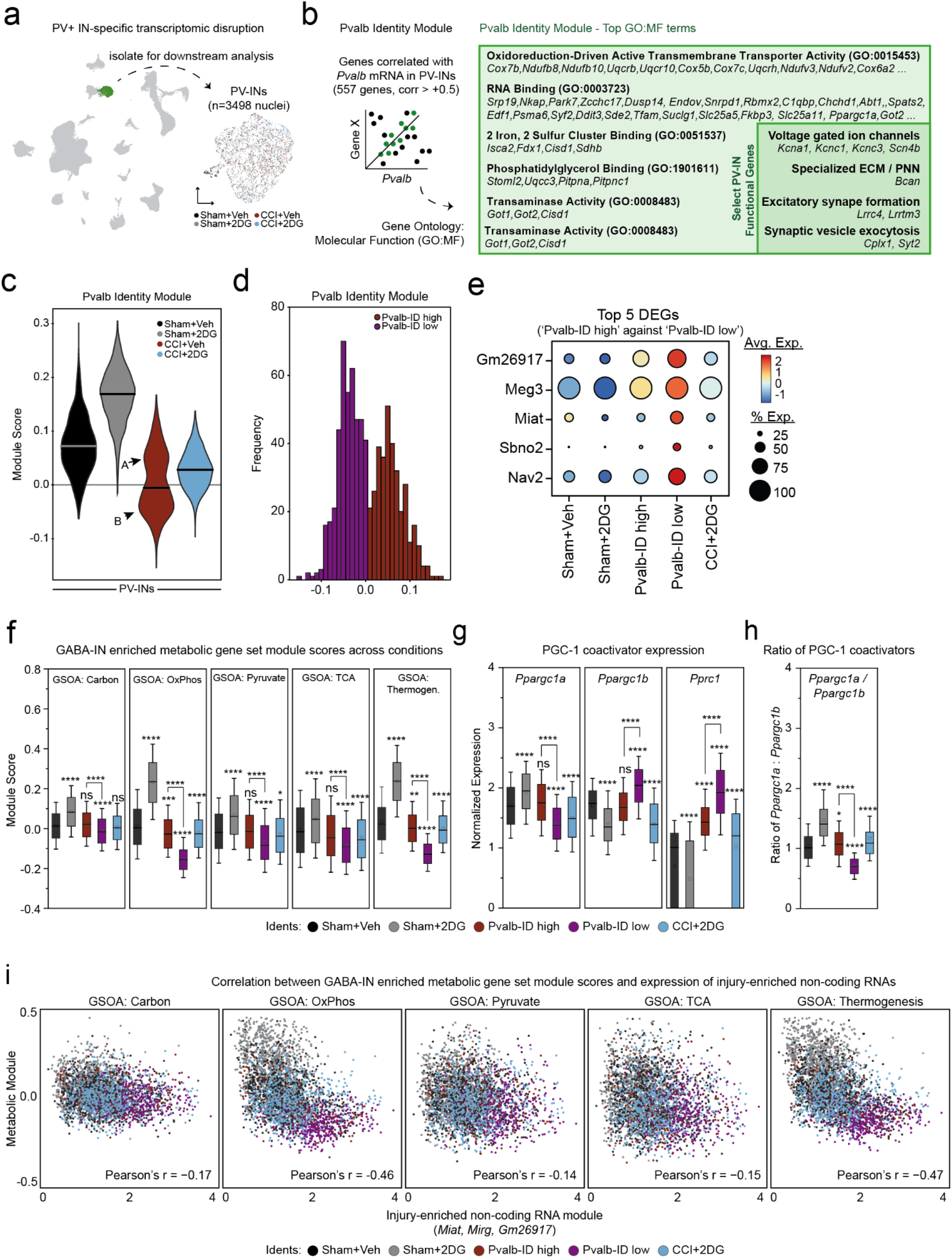
Heterogenous PV-IN populations in CCI+Veh demonstrate relationship between PV-IN identity, oxidative metabolism, and injury associated long non-coding RNAs. **a.** UMAP depiction of all cells (left) and UMAP of isolated PV-INs (right). **b.** Top GO:MF terms for Pvalb Identity Module genes based on overlap; n = 557 genes, positive correlation > 0.5 by Pearson’s correlation. **c.** Violin plot of Pvalb Identity module scores from PV-INs in each experimental condition. ‘A’ and ‘B’ arrows point to two sub-populations of PV-INs in CCI+Veh. **c.** Histogram of Pvalb Identity scores from PV-INs in CCI+Veh. Purple = Pvalb-ID low, PV-INs from CCI+Veh B population; Red = Pvalb-ID high, PV-INs from CCI+Veh A population. **e.** Dotplot depiction of the top 5 enriched genes in Pvalb-ID low PV-INs. Each gene is significantly enriched in Pvalb-ID low group; p < 0.0001 by Wilcoxon Rank Sum with Bonferroni correction. **f.** Boxplots of the module scores for each GSOA term across each PV-IN group; inner squares represent mean values, inner black lines represent median values. Lower and upper box limits represent 25th and 75th percentile, respectively. Error bars represent 10th and 90th percentile. * p < 0.05, ** p < 0.01, *** p < 0.001, **** p < 0.0001 by Wilcoxon Rank Sum with Bonferroni correction **g.** Boxplots showing normalized expression of *Ppargc1a*, *Ppargc1b*, and *Pprc1* in each PV-IN group. **h.** Boxplot showing the ratio of *Ppargc1a:Ppargc1b* mRNA expression in each PV-IN group. * p < 0.05, **** p < 0.0001 by Wilcoxon Rank Sum with Bonferroni correction **i.** Correlation between GSOA module scores and ‘injury-enriched non-coding RNA’ module scores in PV-INs.

While there are limited studies on the role of these lncRNAs in neurons, *Miat*, which was selectively expressed in Pvalb-ID low PV-INs, negatively regulates PPAR signaling and *Cpt1* expression, a target of PGC-1α^58^. With this in mind, we suspected that increased expression of these lncRNAs in injured PV-INs may represent an opposing transcriptional program associated with decreased metabolic capacity and loss of *Ppargc1a*. To that end, we generated module scores based on previous metabolic GSOA terms for PV-INs in each condition (Fig. 6f). As hypothesized, Pvalb-ID low PV-INs had significantly lower metabolic module scores across all GSOA terms compared to Pvalb-ID high PV-INs (Fig. 6f). This was greatest for OxPhos and thermogenesis modules (Fig. 6f). We next assessed whether the PV-IN-enriched PGC-1 coactivators, such as *Ppargc1a*, followed a similar trend. *Ppargc1a* was significantly downregulated in Pvalb-ID low PV-INs, but its levels were unchanged in Pvalb-ID high PV-INs, relative to Sham+Veh (Fig. 6g). Other PGC-1 coactivators (*Ppargc1b* and *Pprc1*) showed similar patterns (Fig. 6g), suggesting large scale disruption of PGC-1 signaling in dysfunctional PV-INs. Lastly, we quantified the correlation between select lncRNAs and metabolic gene expression in PV-INs by creating a gene set module composed of *Miat*, *Mirg*, and *Gm26917* and found a large negative correlation with OxPhos and thermogenesis (Fig. 6i). This suggests a novel relationship between PV-identity, PGC-coactivating molecules, lncRNAs and metabolic gene expression, and suggest opposing transcriptional programs implicated in PV-identity and function.

### Human TBI snRNAseq data show nearly identical metabolic specialization and injury-induced changes in PV-INs

We next performed similar analysis on published human snRNAseq data^23^ from TBI patients and post-mortem control samples. All cortical brain samples were collected from adult individuals (n=12 TBI samples, age range: 22-74; n=5 control samples, age range: 69-87) (Fig. 7a). We isolated 9590 neuronal nuclei (*RBFOX1*, *RBFOX3*) and annotated subclusters (Fig. 7a-b) including a Glut-PN population (*SLC17A7*, *GLS*) and GABA-IN subtype populations (Fig 7b). In control samples, we first validated cross-species PV-IN transcriptional identity and metabolic specialization. Excitingly, we found that the Pvalb-ID module we generated using mouse data robustly identified human PV-INs, and PV-INs were enriched for GSOA metabolic modules compared to other similar neuronal subtypes (Fig. 7c-d), suggesting conserved PV-IN transcriptional identity and metabolic specialization across species. Next, we stratified human PV-INs by injury status and time post injury (Fig 7e). TBI caused a 45% reduction in the median Pvalb-ID score, compared to control samples (p < 0.0001 by Wilcoxon rank sum with Bonferroni correction), and was significantly reduced in TBI samples collected 1-4 days after TBI, relative to those collected 0-1 day after injury (p < 0.0001 by Wilcoxon rank sum with Bonferroni correction) (Fig. 7e). TBI also significantly decreased GSOA term module scores for OxPhos, pyruvate, and thermogenesis (Fig. 7f). Mechanistically, we also found that the median expression of *PPARGC1A* and select interacting TFs implicated in GABA-IN metabolism (*PPARG*, *ESRRG*, *NCOA2*) were significantly decreased in PV-INs after TBI (log_2_FC < −0.1, p < 0.0001 by Wilcoxon rank sum with Bonferroni correction) (Fig. 7g). PGC-1a module expression (*PPARGC1A*, *PPARG*, *ESRRG*, and *NCOA2*) was dramatically reduced by TBI, with a significant decrease in median score from 0-1d to 1-4d and 4-8d (Fig. 7h). Lastly, we examined the expression of lncRNAs found in Pvalb-ID low PV-INs in mice. While only *MIAT* was detected in the human data, its expression was significantly increased in PV-INs after TBI (+1.34 FC, p < 0.0001 by Wilcoxon rank sum with Bonferroni correction) (Fig. 7i). These analyses identify nearly identical findings in human and mouse TBI demonstrating association of PV-IN transcriptional identity, metabolic gene specialization, and opposing regulation of *PPARGC1A* and *MIAT* in PV-INs after TBI. Remarkably, similar re-analysis of neurons from two additional human snRNAseq datasets, human temporal lobe epilepsy^59^ and human schizophrenia^60^, also showed significant reductions in Pvalb-ID scores across PV-INs, suggesting that our Pvalb-ID module detects PV transcriptional dysfunction outside of the TBI context (Supp. Fig. 9).

**Figure 7.**
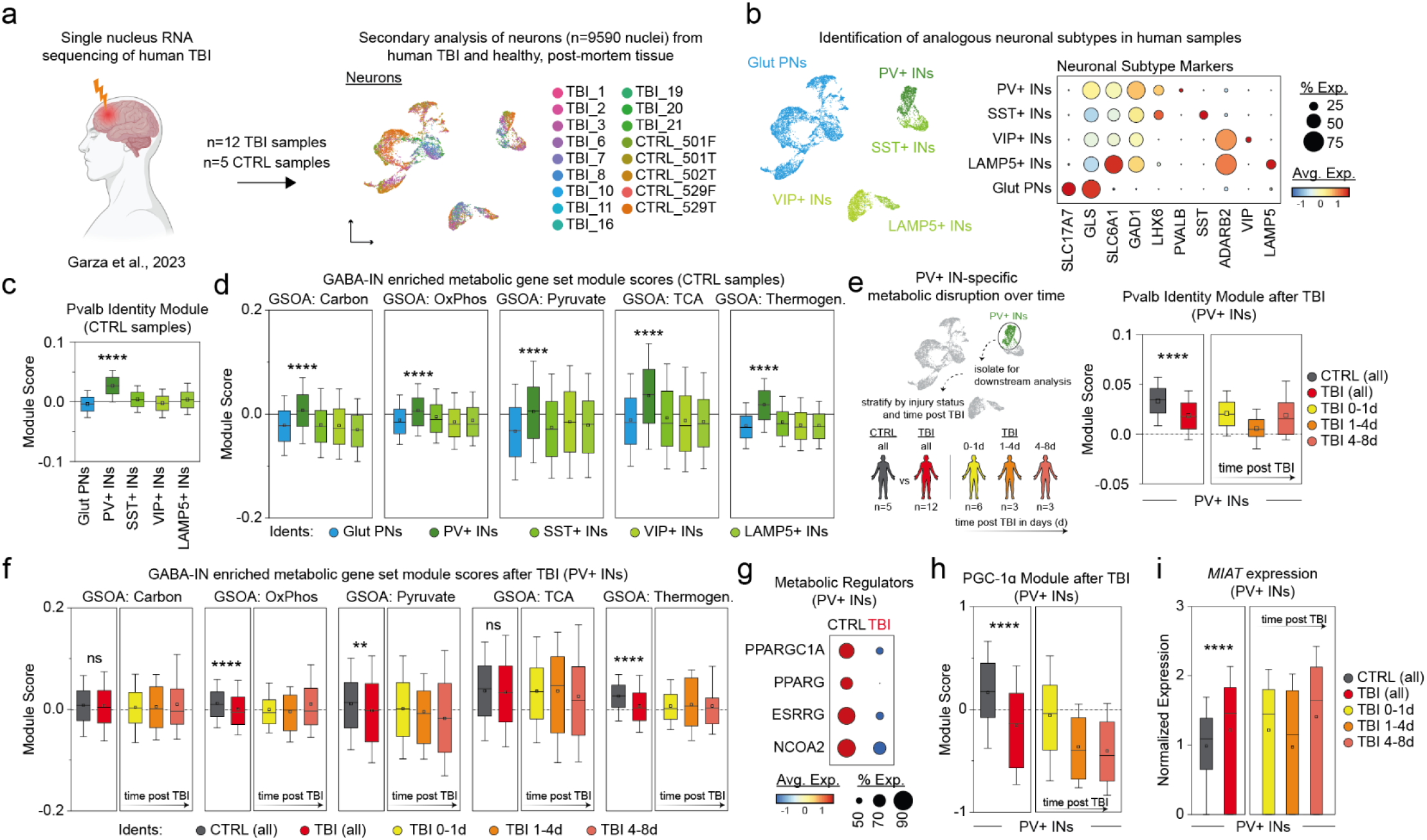
Single nucleus RNAseq analysis of human neurons after TBI confirms PV-IN-specific metabolic enrichment and shows progressive loss of *Ppargc1a* and interacting molecules. **a.** Schematic depiction (*left*) and UMAP depiction (*right*) of reanalyzed neuronal nuclei from published dataset using human TBI and post-mortem CTRL (no TBI) cortical tissue. Nuclei are colored by patient sample **b.** UMAP depiction of human neurons colored by cell type (*left*). Dotplot of neuronal subtype markers (*right*). Values represent average expression. **c**. Boxplot of Pvalb identity module scores across neurons in the CTRL samples showing significant enrichment of Pvalb Identity in PV-INs compared to all other neuron groups. **** p < 0.0001 by Wilcoxon Rank Sum with Bonferroni correction. **d.** Boxplot of module scores for each GSOA term across neurons in the CTRL samples. **** p < 0.0001 by Wilcoxon Rank Sum with Bonferroni correction. **e.** Schematic depiction of the workflow to assess PV-IN-specific metabolic disruption (*left*). Boxplot of Pvalb Identity module scores across PV+INs in CTRL and TBI samples stratified by timepoint (*right*). **** p < 0.0001 by Wilcoxon Rank Sum with Bonferroni. **f.** Boxplot of module scores for each GSOA term across PV+INs in CTRL and TBI samples stratified by timepoint, comparing TBI PV+INs to CTRL PV+INs. ** p < 0.01, **** p < 0.0001 by Wilcoxon Rank Sum with Bonferroni correction. **g.** Dotplot of average expression of *Ppargc1a* and interacting molecules predicted in Figure 4. Values represent average expression across PV+INs in either CTRL or TBI samples. p < 0.0001 by Wilcoxon Rank Sum with Bonferroni correction. **h.** Boxplot of PGC-1α module scores across PV+INs in CTRL and TBI samples stratified by timepoint. PGC-1α module is comprised of *PPARGC1A, PPARG, ESRRG,* and *NCOA2*. **** p < 0.0001 by Wilcoxon Rank Sum with Bonferroni correction. **i.** Boxplot of *MIAT* mRNA expression in PV+ INs across CTRL and TBI samples stratified by timepoint. **** p < 0.0001 Wilcoxon Rank Sum with Bonferroni correction.

## Discussion

Neurometabolic dysfunction occurs in many neurological and psychiatric disorders including TBI, epilepsy, Alzheimer’s disease, schizophrenia, and more^47,61–66^. Meanwhile, PV-INs, the largest class of GABAergic interneurons^27^, are thought to be uniquely susceptible to metabolic pressure and are commonly lost and/or dysfunctional across many of these disorders^1–7^. In TBI there is a well-defined neurometabolic cascade that leads to neuronal and circuit dysfunction, and PV-IN loss^65,66^. How metabolic dysfunction associated with TBI, and other disorders, might underlie PV-IN-specific dysfunction, and the molecular differences that drive this cell type-specific dysfunction are not established. Here, we used snRNAseq in male mouse cortex 3 days after CCI to show that PV-INs are uniquely specialized in their expression of multiple metabolic pathways (OxPhos, TCA, thermogenesis, carbon and pyruvate metabolism), that specialization is lost after CCI, and in vivo treatment with 2DG, which protects PV-INs and inhibitory network function after injury^13^, restores PV-IN metabolic specialization and transcriptional identity. By examining upstream regulators of metabolic genes enriched in GABA-INs, we identified *Ppargc1a* as a PV-IN-selective regulator of injury-induced transcriptional changes. Since both TBI and metabolic dysfunction of PV-INs are associated with loss of PNN^12,24,51^, we examined the expression of PNN-related CSPGs. We found that brevican expression was decreased in astrocytes and PV-INs by CCI and partially restored by 2DG, and that the amount of PNN labeling followed a similar pattern. We also observed that the ratio of PV-INs surrounded by PNN was reduced by CCI and increased by 2DG treatment. We generated a unique transcriptional module that correlates with *Pvalb* expression (gene encoding PV protein) in PV-INs and can be used to quantify the robustness of PV-IN transcriptional identity. The overall expression of this Pvalb-ID module is decreased after TBI, but is restored by 2DG treatment, and includes genes associated with metabolic pathways enriched in GABA-INs (*Me3, Dlat, Cox6a2*), ion channels that mediate fast-spiking (*Kcna1, Kcnc1*), specializations of inhibitory synaptic function (*Cplx1, Syt2*), and transcriptional regulators of mitochondrial function and biogenesis (*Tfam, Ppargc1a*).

Using the PV-identity module score, we report a bifurcation in PV-IN transcriptional identity after injury and identify lncRNAs associated with inflammation and cellular stress^67,68^ (*Miat* and *Meg3*) that are increased in Pvalb-ID low cells. The abundance of these lncRNAs inversely correlates with PV-IN metabolic specialization, suggesting they may be part of a transcriptional program that opposes PGC-1α signaling downstream to reduce PV-IN specialization. Finally, we applied the same analysis to a published human TBI snRNAseq dataset and found that PV-IN metabolic specialization is seen in humans, loss of that specialization occurs in patients after TBI, and PGC-1α-related TFs are decreased while *MIAT* is increased in human PV-INs after TBI. These changes appear to be progressive over the first post-injury week in human patients. Together, this provides new tools to quantify PV-IN transcriptional specialization, delineates key metabolic pathways essential to PV-IN function that are disrupted by TBI but restored by 2DG, and identifies transcriptional co-activators and lncRNAs in maintaining and disrupting PV-IN transcriptional specialization, respectively. The striking similarities between the rodent and human TBI data suggest metabolic interventions after TBI, and particularly, the targeting of select molecules (*PPARGC1A*, *MIAT*) identified in Pvalb-ID low PV-INs, may have translational potential to improve inhibitory function after TBI or act as biomarkers for TBI severity.

While glia are known to be metabolically specialized^69,70^, whether different neuronal subtypes have specialized metabolic systems to support their diverse functions is not well-known. PV-INs, thought to have higher metabolic demands to support fast spiking, are known to be enriched in mitochondria^71^ and express unique metabolic enzymes^12^ (Cox6a2) that when lost compromise their function. PGC-1α, a transcriptional co-activator that interacts with PPAR-γ and other co-activators to shape metabolic activity across cell types and tissues, has previously been identified as critical in coordinating metabolic and synaptic function of PV-INs^45–47^. Our results confirm these findings and demonstrate that these systems are disrupted after TBI. Whether similar changes occur in other pathologies is unknown, but multiple studies link metabolism to other PV-IN specializations including maintenance of PNN^12,51^, fast-spiking APs, and the fidelity of presynaptic function during heightened activity^46^. Underscoring the importance of studying cell type-specific neuronal metabolism, we find that GC-MS metabolomic analysis replicates known TBI-induced changes but fails to capture significant changes in metabolic state caused by 2DG (Supp Fig 3a-b). While the transcriptional changes we report in PV-INs after 2DG treatment are striking, PV-INs make up a minority of neurons in the cortex and thus may undergo large changes that cannot be detected with standard metabolomic approaches. Additionally, it may also suggest that 2DG treatment drives PV-INs to use different metabolic systems that allow them to function and survive in a compromised metabolic milieu. We demonstrate that ubiquitous changes to metabolic pathways (e.g. indiscriminate downregulation of OxPhos and mitochondrial function) can have cell type-specific downstream consequences through cell type-specific interacting molecules (e.g. PV-IN-specific interaction with PGC-1α).

A caveat of this study is that metabolic gene expression is subject to post-transcriptional and feedback regulation meaning transcriptional changes are only part of metabolic regulation^72^. Additionally, despite the abundance of nuclear-encoded mitochondrial genes, single *nucleus* RNAseq does not typically capture mitochondrially encoded genes. Finally, our animal studies focus only on males, and the human data we use is underpowered to examine sex-specific differences in PV-IN biology or TBI-associated metabolic response, which is known to be different in males versus females^73^. Those caveats aside, our study supports the utility of single cell transcriptomics for identifying cell type-specific metabolic genes sets that are physiologically and therapeutically relevant.

The use of metabolic therapies to treat neurological disorders is not new, but how these therapies affect brain function is not well understood. The ketogenic diet, which shifts the brain from reliance on glycolysis to ketosis^74^, has a long history as an anticonvulsant and neuroprotective therapy^75^, and provided the rationale to use 2DG to inhibit glycolysis after injury^76^. 2DG is known to acutely reduce neuronal activity^77,78^, activate the NRSF pathway via changes in NADH/CtBP^79^, promote the adaptive ER stress-ATF4 pathway^36^, and enhance BDNF expression^77^. Our data supports that 2DG enhances oxidative metabolism, particularly in GABA-INs, in parallel to its other effects. We suspect that 2DG’s effects on the metabolic transcriptome are mediated by inhibition of glycolysis, altering intracellular metabolite levels (e.g. NADH, FAD, acetyl-coA) to increase the expression of oxidative metabolism genes. Because PV-INs are enriched in mitochondria and are thought to be more reliant on oxidative phosphorylation compared to other neuronal subtypes, 2DG’s effects may be particularly effective in promoting PV-IN health. Similarly, because PGC-1α promotes mitochondrial biogenesis^80^, its upregulation specifically in GABA-INs underscores the intimate link between oxidative phosphorylation and inhibitory circuit function, as well as 2DG’s ability to protect PV-INs after injury. Previous work on PGC-1α in PV-INs demonstrates that *Pvalb* and *Syt2* expression are downstream of PGC-1α and link its expression to PV-IN transcriptional health and function^45,47^. Similarly, studies in muscle and cancer cells demonstrate that increases in PGC-1α increase mitochondrial biogenesis, oxidative phosphorylation activity, and resistance to metabolic challenge^81,82^. Taken together, this suggests that restoration or enhancement of PGC-1α in PV-INs may restore PV-IN identity genes, including oxidative metabolism gene sets, and preserve their specialized function following TBI.

In addition to genes already implicated in PV-IN function, our study also identifies novel targets that are dysregulated after TBI. Multiple lncRNAs were increased in Pvalb-ID low PV-INs, and their abundances inversely correlate with metabolic specialization. *MIAT*, originally identified as a risk factor for myocardial infarction in clinical populations, is associated with inflammatory signaling^68^, promotes apoptosis after hypoxia^83^, and is a negative regulator of PPARα signaling^58^. *MEG3* participates in epigenetic regulation of transcriptional control and promotes cell death after ischemia^84^. Interestingly, *MIAT* is elevated in PV-INs in the brains of schizophrenia patients and is increased in blood exosomes^85^, suggesting it may have value as a biomarker of PV-IN dysfunction after TBI. These novel links between lncRNAs, TBI, and metabolism suggest complex cell type-specific interactions that shape PV-IN identity and function.

Finally, re-analysis of a published snRNAseq dataset^23^ of human TBI patients show nearly identical PV-IN metabolic changes as we report in mice. The PV-identity module developed using data from mice robustly identifies human PV-INs and is decreased following TBI. PV-INs are enriched in multiple metabolic pathways compared to other neuronal subtypes, and *Ppargc1a* is decreased while *MIAT* expression is increased in PV-INs after TBI. Human data also supports dysregulation of additional PGC-1α-interacting TFs in PV-INs after TBI (*ESRRG*, *PPARG*, and *HNF4A*) that were below detection in mice for quantitative analysis. Remaining questions include whether similar metabolic and signaling systems disrupt PV-IN function in other pathologies, whether Pvalb-ID low PV-INs remain into the chronic phase of TBI or are destined for cell death, and how changes in glial function (reactivity, inflammatory signaling, metabolic support) contribute to PV-IN metabolism and dysfunction. Excitingly, human snRNAseq and human bulk RNAseq studies of other neurological disorders suggest similar mechanisms might be at play. In human temporal lobe epilepsy, PV-INs show decreased Pvalb identity, as well as decreased *PVALB*, *PPARGC1A*, and *ESRRG* expression^59^. In human AD, *PVALB* is decreased in PV-INs at all stages of disease progression, while *PPARGC1A* is significantly decreased late in disease progression^86^. In a bulk RNAseq study from human chronic traumatic encephalopathy (CTE), *PVALB*, *PPARGC1A*, *ESRRG*, and *PPARG* are all significantly decreased^87^. As tools develop to study single cell metabolomics, and we better understand the dynamic metabolic interactions that control cellular and circuit function in health and disease, we hope to leverage cell type-specific metabolic differences to reduce the loss of metabolically sensitive PV-INs, and other cell types, across neurological conditions.

## Methods

### Animals

All animal procedures were performed in accordance with the Tufts University School of Medicine’s Institutional Animal Care and Use Committee. All experiments were performed on adult C57BL/6 male mice. Mice were obtained from Charles River Laboratories, Jackson Laboratories, or bred in-house. All mice used for snRNAseq experiments were obtained from Charles River Laboratories. Animals were kept on a standard 12-hour light/12-hour dark cycle and fed ad libitum with regular chow diet and water.

### CCI

TBI was modelled using CCI, as previously described^13^. Mice were anesthetized using inhaled isoflurane in oxygen (4% for induction, 2% for maintenance). Once head-fixed in a stereotaxic frame, the scalp was sterilized, and a vertical midline incision was made. A 5-mm craniectomy was performed left of the midline, between bregma and lambda. The resulting skull flap was removed. The impact was made using Leica Benchmark Stereotaxic Impactor, with a 3-mm diameter piston, 3.5 m/s velocity, 400-ms dwell time, and 1-mm depth. Following the impact, sutures were used to close the scalp incision. The skull flap was not re-inserted. Sham animals were anesthetized for the same approximate time but did not receive an impact or craniectomy. Animals were all single housed following surgery until the day of sacrifice.

### In vivo 2DG treatment

Animals were injected intraperitoneally with 200µL of 2DG dissolved in sterile saline at a dose of 250 mg/kg. Vehicle injections were 200µL injections of sterile saline. The first injection was administered approximately 20 minutes after CCI or sham surgery. Daily injections were then continued for 3 days until the time of sacrifice.

### Immunohistochemistry

Animals were transcardially perfused with 50 mL of chilled PBS, followed by 50mL of 4% paraformaldehyde (PFA) in 0.1 M phosphate buffer. Brains were harvested and placed overnight in 4% PFC solution at 4°C, followed by 3 days in a 30% sucrose solution. Coronal slices (40 μm) were made on a Thermo Fisher Scientific Microm HM 525 cryostat. Slices were washed with PBS and then incubated in blocking buffer (10% normal goat serum [NGS], 5% bovine serum albumin [BSA] in PBS with 0.2% Triton X-100 ((PBS-T)) at room temperature for 1 hour. The slices were incubated overnight at 4°C with primary antibody in PBS-T with 5% NGS/1% BSA. A primary antibody against PV (mouse; Swant PV235; 1:1000) was used in addition to *Wisteria floribunda* Agglutinin (WFA) (Sigma-Aldrich, 1:200). Slices were washed with PBS and incubated with secondary antibody (anti-mouse Alexa 647, Alexa 488-conjugated streptavidin) in PBS-T with 5% NGS/1% BSA for 2 hours at room temperature. Lastly, slices were rinsed with PBS and DAPI, then mounted. PV and PNN images were taken on a Keyence epifluorescence microscope using a 10x objective. For each experiment, identical laser and capture parameters were used across all experimental groups.

### Tissue harvest and nuclei isolation for snRNAseq

Mice were anesthetized in an isoflurane chamber and rapidly decapitated. Brains were quickly harvested and placed in a chilled slicing solution (in mM: 234 sucrose, 11 glucose, 24 NaHCO3, 2.5 KCl, 1.25 NaH2PO4, 10 MgSO4, 0.5 CaCl2) equilibrated with 95% O2/5% CO2. The brain was mounted to a Leica VT1200S vibratome, and the slicing chamber was filled with chilled slicing solution. 1mm thick coronal slices were taken and quickly placed into a petri dish containing chilled Hibernate-A media (ThermoFisher, Cat No. A1247501), and perilesional cortical tissue ipsilateral to the site of injury was manually dissected under a dissection microscope, with care to not include underlying white matter. Tissue from a corresponding location was used from Sham animals. The dissected cortical tissue was placed into a pre-chilled dounce homogenizer on ice, along with 4mL of lysis buffer (Millipore Sigma, Product No. 08168). The tissue was homogenized then transferred to a 15mL conical tube on ice and incubated for 5min, stirring occasionally. After incubation, the sample was centrifuged at 500rcf for 6min at 4°C. The supernatant was removed, and the pellet was then resuspended in lysis buffer and incubated on ice for 5 minutes. Following incubation, the sample was passed through a 30µm strainer (Miltenyi Biotec) and centrifuged at 500rcf for 6min at 4°C. After, the supernatant was removed, and the pellet was resuspended in resuspension buffer (2%BSA in 1XPBS containing 0.2 U/µL RNase inhibitor) and centrifuged again at 500rcf for 6mins at 4°C. The supernatant was removed, and the pellet was resuspended once more in 500 µL of resuspension buffer. 10µL of nuclei suspension diluted 1:1 in trypan blue was loaded into a hemocytometer, the concentration of nuclei was determined, and resuspension buffer was added as needed to reach a final concentration of ∼1000 - 1200 nuclei/µL.

### 10x Chromium single nuclei capture of cortical cells

Isolated cortical nuclei [resuspension buffer: 2%BSA in 1XPBS containing 0.2 U/µL RNase inhibitor] were loaded into the 10x Chromium chip for subsequent preparation using the 10x Genomics Single Cell 3′ v3 and v3.1 chemistry according to the manufacturer’s instructions. Approximately 8,000 nuclei were loaded at a nuclear concentration of ∼1000 - 1200 nuclei/µL. For cDNA amplification PCR, 11 total cycles were applied. cDNA library quality and concentration were analyzed on an Agilent Fragment Analyzer. Final cDNA libraries were sequenced on a NextSeq 550 using High Output V2.5 chemistry.

### Genome alignment and quality control (QC) processing of single cell transcriptomes

The raw sequence data was used as input for cellranger mkfastq to create the fastq files, followed by cellranger count for read alignment to mm10 mouse genome and generation of output matrices [CellRanger v3.1; 10x Genomics]. The filtered feature barcode matrix was then used for downstream analysis in R-Seurat v4^88^. Using R-Seurat, for QC, we removed genes that were not present in at least 5 cells, and cells that did not have a minimum of 350 genes. Additional QC was to remove cells that had greater than 6% of their library comprised of mitochondrial genes. Suspected doublets were removed by plotting the number of UMIs against the number of genes and removing cells that were outliers.

### CCA integration, clustering, and cell type identification

To improve our cell type identification across different biological conditions, we performed CCA integration across all our single cell transcriptomic datasets^25^. This process computed 2000 integrated genes that were then used as input for PCA. A UMAP was then run using the first 50 principal components of PCA. This was used to project our single cell transcriptomes in a single UMAP space, as seen in Figure 2a.

For clustering, we used the integrated features as input and utilized the default graph-based clustering approach described in Seurat v3. For the ‘FindNeighbors’ function, we used principal components 1:50, and the remaining default arguments. For ‘FindClusters,’ we used a resolution of 1, and the remaining default arguments. This resulted in over-clustering of the data (i.e., there were more clusters than biological cell types). To resolve each cluster into its biological cell type identity, we performed DE analysis that allowed us to determine the most highly enriched genes (p adjusted < 0.05) for each cluster relative to the remaining clusters. We then iteratively assigned each cluster based on their enriched and selective gene expression using established publicly available transcriptomic datasets^89,90^.

### Pathway analysis using gene set enrichment analysis

To elucidate CCI-, and 2DG-associated pathways, we first generated lists of differentially expressed genes (DEGs; inclusion criteria: min.pct > 0.1, absolute log_2_FC > 0.25, p < 0.1 by Wilcoxon rank sum with Bonferroni correction) using the ‘FindMarkers’ function within Seurat for each cell type that we identified across each comparison. Sham+Veh versus CCI+Veh produced the CCI-associated DEGs. Sham+Veh versus Sham+2DG produced the 2DG-healthy-associated DEGs. CCI+Veh versus CCI+2DG produced the 2DG-CCI-associated DEGs. Respective DEG lists (p adjusted < 0.1, min.cells = 10% of cell type) were then used as input into Ingenuity Pathway Analysis (IPA) software for Canonical Pathway Analysis^30^. The top 25 affected pathways (p < 0.05) were determined in decreasing order of significance.

### Neuronal metabolic gene expression enrichment analysis

The list of metabolic genes that was used in the metabolic gene expression analysis in Figure 4 was obtained from https://www.genome.jp/kegg/pathway.html. Initial DE analysis (DEGs; inclusion criteria: absolute log_2_FC > 0, p < 0.05 by Wilcoxon rank sum with Bonferroni correction) of metabolic gene expression compared the average expression of each gene across 3 groups: neurons (L2/3 Glut PNs, L4 Glut PNs, L5/6 Glut PNs, PV-INs, SST-INs, VIP-INs, and LAMP5-INs), glia (astrocytes, oligodendrocytes, OPCs 1, and microglia), and vasculature (endothelial cells, pericytes, and VLMC). The DE comparisons were as follows: neuron-enriched (neuron vs all), glia-enriched (glia vs all), and vasculature-enriched (vasculature vs all). The top 100 neuron enriched DEGs were determined in order of decreasing p value.

To determine the distribution of expression of the neuron enriched metabolic genes, a second iteration of DE analysis was performed, using the same inclusion criteria as before. Now, DE analysis was performed on the neuron enriched metabolic genes, comparing the average expression of neuron enriched metabolic genes in Glut PNs (L2/3 Glut PNs, L4 Glut PNs, L5/6 Glut PNs) to the average expression of those genes in GABA-INs (PV-INs, SST-INs, VIP-INs, LAMP5-INs). Neuron enriched metabolic genes were determined to be either Glut PN enriched, GABA-IN enriched, or neither (p adjusted > 0.05). The same neuron enriched genes were used as input across all experimental conditions in Figure 4c.

Gene set overrepresentation analysis (GSOA) was performed by comparing the significantly enriched metabolic genes found in GABA-INs or Glut PNs against a background list of all neuron-enriched metabolic genes to determine the overrepresented pathways in GABA-INs and Glut PNs^55–57^. The pathway database used was KEGG.

### GABA-IN metabolic enrichment correlation analysis

To evaluate whether the increase in number of enriched metabolic genes observed in CCI+2DG represented a restoration back to Sham+Veh levels (i.e., the same enriched genes in Sham+Veh are the same enriched genes in CCI+2DG), we took the neuron enriched metabolic genes from Sham+Veh and plotted the distribution of those genes along the x-axis based on the enrichment level observed in Sham+Veh. Along the y-axis, each gene was plotted based on its enrichment level observed in each respective condition. This generated 4 scatter plots, in which a linear fit of slope = 1 would represent a restoration to Sham+Veh metabolic enrichment levels. A linear fit was then applied, and a Pearson correlation was applied to evaluate the fit of the data.

### Upstream regulator analysis

To identify potential upstream regulators of metabolic enrichment in GABA-INs, we took the list of GABA-IN enriched metabolic genes and performed GSOA as previously described, however, the pathway database was set to “TF.Target.TF.Perturbations.Followed.by.Expression” which includes target gene sets from transcription factor (TF) perturbation experiments^55^. This generated a list of upstream TFs that were experimentally observed to regulate a significantly overlapping set of genes with our GABA-IN enriched metabolic gene set. We then cross-referenced that list of TFs against the PV-IN CCI-associated DEGs and PV-IN 2DG-CCI-associated DEG list to identify key DEGs from our dataset that coincide with the implicated pathways from our GSOA analysis. Prioritization was based on the significance of overlap with the predicted TF, the level of cell type-specificity of the pathway in question, and whether the candidate was observed to be regulated by both CCI and 2DG treatment. Additional consideration was also given to whether the candidate had been previously implicated in injury, implicated in PV-IN biology previously, and whether the candidate was suspected or demonstrated to also play a role in PV-IN neurotransmission.

### Protein-protein interaction network analysis

To determine predicted protein-protein interactions between predicted upstream regulators of GABA-IN-enriched metabolic genes, we started with the list of molecules (TFs) from our list of predicted upstream regulators in Figure 4f. Next, we filtered the list of molecules by false discovery rate (FDR) and normalized enrichment score (NES) (FDR < 0.05, NES < 0.01) and submitted the filtered list to the STRING database^42^. K-means clustering was applied (k=10) and we selected our cluster of nodes of interest based on annotated functional association related to oxidative metabolism and mitochondrial function.

### Tissue prep for GC-MS

To assay relative levels of cortical metabolite concentrations, we performed gas chromatography mass spectrometry (GC-MS) on resected cortical brain tissue from Sham and CCI animals. 3 days following surgery, mice were cervically dislocated and brains were quickly removed. Once the brain was removed, it was placed in a small dish containing ice-cold 1XPBS. Under a dissection microscope, the perilesional cortex was harvested; similar cortical regions were collected from Sham brains. The resected tissue was then quicky washed in ice-cold double-distilled water twice, before being pat dried, and placed in a weigh boat floating in liquid nitrogen. Once the tissue was completely frozen (60-120 seconds), the tissue was placed in a −80°C freezer until GS-MS. Care was taken to work quickly. Each harvest was completed within 5 minutes, from decapitation to storage at −80°C.

### GC-MS: Metabolite Extraction

We conducted metabolite extraction for GC-MS analysis by subjecting samples to sonication (3 pulses for 10 seconds at an amplitude of 5 with 5 second pauses between each pulse) in a 1:1 methanol:water solution. After extraction, we centrifuged the samples at 4°C for 10 minutes at 15,000 rpm, leading to the separation of the insoluble pellet (protein fraction) and the supernatant (polar fraction). We promptly removed the supernatant, snap-froze it using liquid nitrogen, and stored it at −80°C until derivatization.

To the protein fraction, we conducted multiple washes, starting with 50% methanol. After each wash, we centrifuged the samples at 4°C for 10 minutes at 15,000 rpm. This cycle was repeated three times, concluding with a last wash using 100% methanol. Subsequently, we removed the supernatant and subjected the pellet to vacuum centrifugation using a CentriVap Benchtop Vacuum Concentrator connected to a CentriVap −105°C Cold Trap (Labconco) for 30 minutes under a pressure of 10^−3^ mBar to achieve drying.

The dried pellet underwent hydrolysis by adding a 3M HCl solution and incubating the samples at 95°C for 2 hours. We terminated the hydrolysis by adding 100% methanol, for a final concentration of 50% methanol. After incubation on ice for 30 minutes, we centrifuged the samples at 4°C for 10 minutes at 15,000 rpm. The supernatant was transferred to a new tube and stored at −80°C until derivatization.

### GC-MS: Sample Derivatization

We derivatized the dried polar and protein samples using methoxyamine HCl (20 mg/mL) in pyridine and N-methyl-trimethylsilyl-trifluoroacetamide (MSTFA; Thermo Fisher Scientific). For the polar fraction, we added 50 μL of methoxyamine HCl and incubated the samples at 30°C for 90 minutes. As for the protein fractions, we added 70 μL of methoxyamine HCl, followed by centrifugation at 15,000 rpm for 10 minutes. We then transferred 50 μL of the supernatant to a v-shaped amber glass chromatography vial. Finally, we added 80 μL of MSTFA to each sample and incubated them at 37°C for 30 minutes.

### GC-MS Quantification

We utilized an Agilent 8890 gas-chromatography (GC) system paired with a 5977C single quadrupole mass spectrometry detector equipped with an InertPlus Electron Ionization (EI) source. The experimental setup followed established procedures (Young et al., 2020). Specifically, 1 μL of derivatized sample was injected into a J&W HP-5ms Ultra Inert GC column (Agilent Technologies, #19091S-433UI) using a 1:10 split mode. Ultra-high purity helium (Airgas #UN1046) flowed at 0.687 mL/min with a pressure of 4.3 psi. The initial temperature was maintained at 60°C for 1 minute, then increased to 325°C at a rate of 10°C/min and held for 10 minutes. EI energy was set at 70 eV, the transfer line temperature at 290°C, and the source temperature adjusted to 250°C for polar and 280°C for non-polar samples. Data acquisition occurred in scan mode (50-550 m/z).

To identify metabolites, we matched EI fragmentation patterns and retention times with the Fiehn 2013 Metabolomics RTL Database (from Agilent) in the Automated Mass Spectral Deconvolution and Identification System (AMDIS) software. Relative abundance was determined using the Data Extraction for Stable Isotope-labeled Metabolites (DEXSI) software, normalized to total protein abundance by summing amino acid signals from the non-polar fraction.

### Single molecule fluorescent *in situ* hybridization (smFISH) using RNAscope

Animals were perfused, and brains were treated as previously described for immunohistochemistry. Coronal brain sections (14 µm) were made on a Thermo Fisher Scientific Microm HM 525 cryostat, and sections were mounted directly onto slides. The tissue slides were then stored at −80°C. smFISH was done according to manufacturer’s directions (ACDbio – RNAscope Multiplex Fluorescent Reagent Kit (Cat No. 323130)), with exception of one minor change. To improve adherence, pre-mounted slides were baked at 60°C for 1 hour prior to the first washing step. The procedure was done to manufacture specifications. The probes used were all from the ACDbio catalog of probes: *Ppargc1a* – C1 (Cat No. 402121), *Sox9* – C2 (Cat No. 401051), and *Gad1* – C3 (Cat No. 400951). Images were taken on a Nikon A1R confocal microscope with a ×40 oil objective (Olympus). Image processing was done using ImageJ^91^.

### Secondary analysis of published human snRNAseq data

Human traumatic brain injury (TBI): Feature-barcode matrices for each patient sample obtained from Garza et al^23^ were downloaded from NCBI GEO under accession GSE209552^23^. R-Seurat was used to create a merged Seurat object containing all human snRNAseq data. QC was performed as described in original study (nuclei with <1000 genes were removed). Cell clusters were determined and annotated as described previously for rodent snRNAseq data. Cell clusters enriched for RBFOX1 and RBFOX3 (Cluster X vs all; log_2_FC > 0.25, p < 0.0001 by Wilcoxon rank sum with Bonferroni correction) were determined to be neurons and subset for downstream analysis. All downstream analysis was performed as previously described for rodent snRNAseq data.

Human temporal lobe epilepsy (TLE): Feature-barcode matrices for each patient sample obtained from Pfisterer et al^59^ were downloaded from EGA under accession EGAS00001002882. R-Seurat was used to create a merged Seurat object containing all sample data. Cell and group annotations were determined in the original publication and were downloaded as metadata along with count matrices. All downstream analysis was performed as previously described for rodent snRNAseq data.

Human schizophrenia (SCZ): Feature-barcode matrices for each patient sample obtained from Batiuk and Tyler et al^60^ were downloaded from Zenodo under DOI: 10.5281/zenodo.6921620. R-Seurat was used to create a merged Seurat object containing all sample data. Cell and group annotations were determined in the original publication and were downloaded as metadata along with count matrices. Non neuronal cells were removed for downstream analysis. All downstream analysis was performed as previously described for rodent snRNAseq data.

## Acknowledgements

We thank members of the Dulla, Sun, and Gentry Labs for helpful comments on the project and manuscript. We thank Mary Sommer for helpful support with immunohistology and FISH. We thank Jacqueline Garcia for helpful support and feedback. We thank Dr. Paul Robson (Jackson Labs) for helpful initial discussions in designing single cell experiments and Dr. Avtar Roopra (University of Wisconsin) for their helpful comments on addressing issues of sampling bias. We thank Dr. Yunhua Zhu for assistance with developing and optimizing nuclei isolation and helpful advice throughout. We thank Drs. Maribel Rios, Jamie Maguire, and Bree Aldridge for helpful support throughout the project on experimental design and statistical analysis. We thank the TUCF Genomics Core, in particular, Irina Grinvald, for assistance with sequencing. This work was supported by NIH (NS098009, TW011225 to CGD; AG066653, AG078702 to RCS; and NS116824 to MSG), HHMI Gilliam Fellowship (SQ), and DoD (W81XWH-18-1-0669 EP170053, W81XWH2210769 EP210014, W81XWH-17-1-0531 EP160021 to CGD), and Sail for Epilepsy (CGD).

## Data Availability

The rodent single-nucleus RNA-sequencing (snRNAseq) data generated in this current study is available for download on a public repository. All human snRNAseq data were obtained from published studies and are available for download on their respective repositories (see ‘Methods’). All other data are available from the corresponding author on reasonable request.

## Code Availability

No custom code was used to generate or analyze data from this current study. All software used is included in the manuscript (see ‘Methods’).

---

**Supplementary Figure 1.**
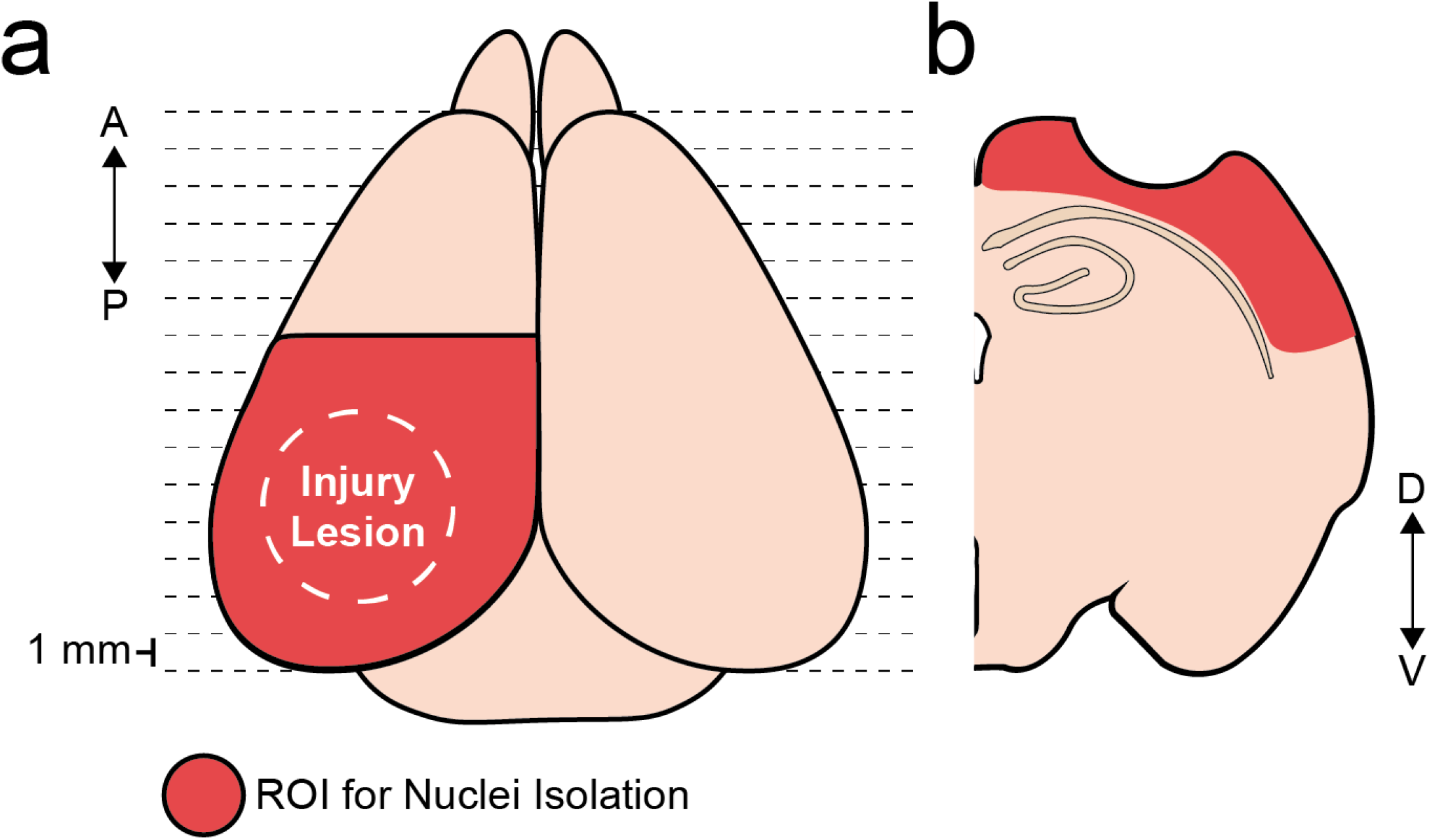
Schematic of cortical tissue dissection for nuclei isolation. **a.** Dorsal view depiction of a mouse brain. Dotted horizontal lines represent where coronal slices were made during tissue dissection process. **b**. Cartoon depiction of a coronal slice generated in (**a**.). Red shaded regions indicate regions of cortical tissue harvested for downstream snRNAseq experiments and GC-MS experiments.

**Supplementary Figure 2.**
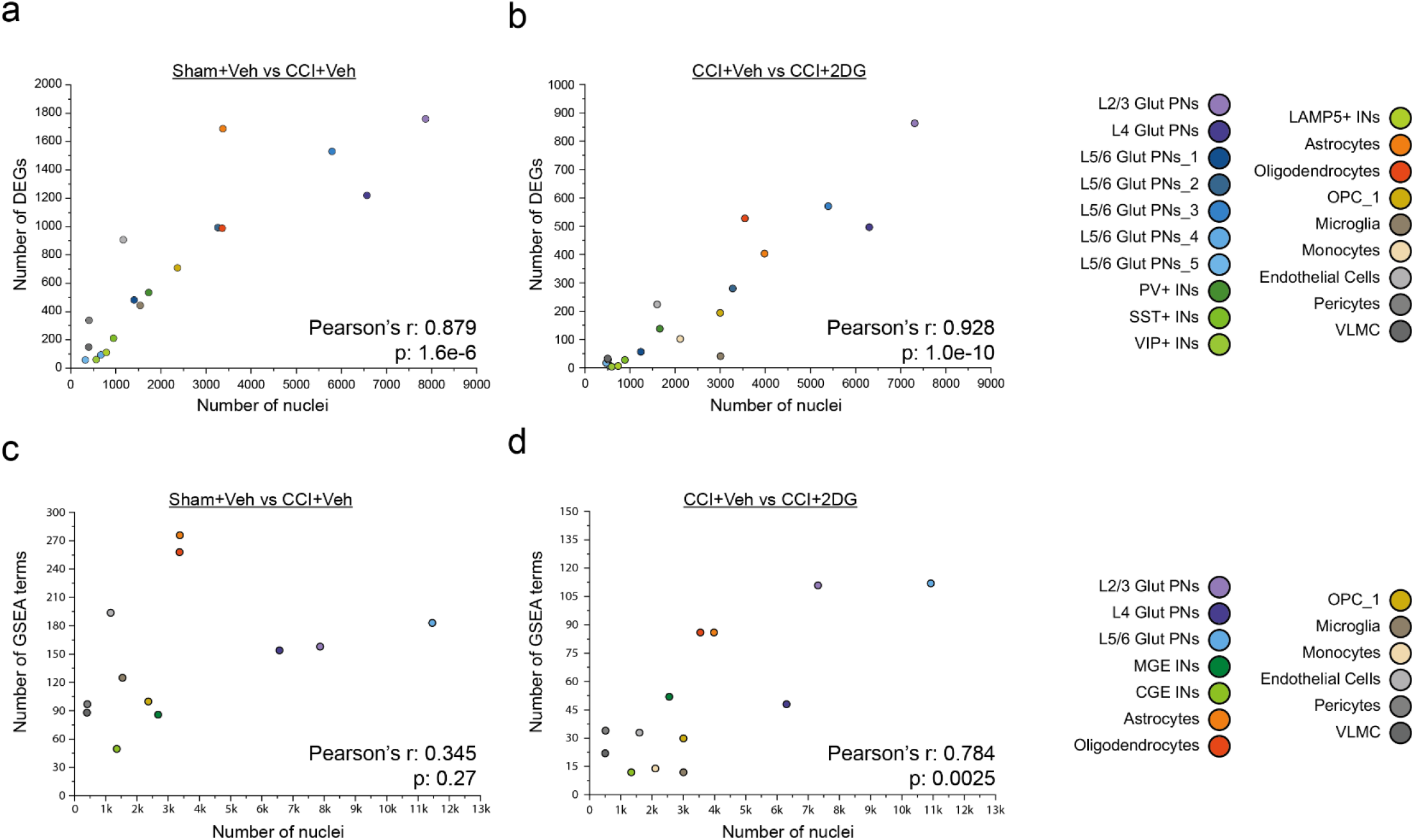
Correlation between number of nuclei and number of DEGs and GSEA terms. **a-b**. Correlation between number of nuclei per cell type (x-axis) and number of DEGs (y-axis). DEGs determined by Wilcoxon Rank Sum with Bonferroni correction (adjusted p < 0.1, expressed in minimum of 10% of cells per cluster. Comparisons: Sham+Veh vs CCI+Veh in (**a**.) and CCI+Veh vs CCI+2DG in (**b**.). **c-d**. Correlation between number of nuclei per cell type (x-axis) and number of GSEA terms (y-axis). GSEA terms determined using IPA software (p < 0.05). Comparisons: Sham+Veh vs CCI+Veh in (**c**) and CCI+Veh vs CCI+2DG in (**d**).

**Supplementary Figure 3.**
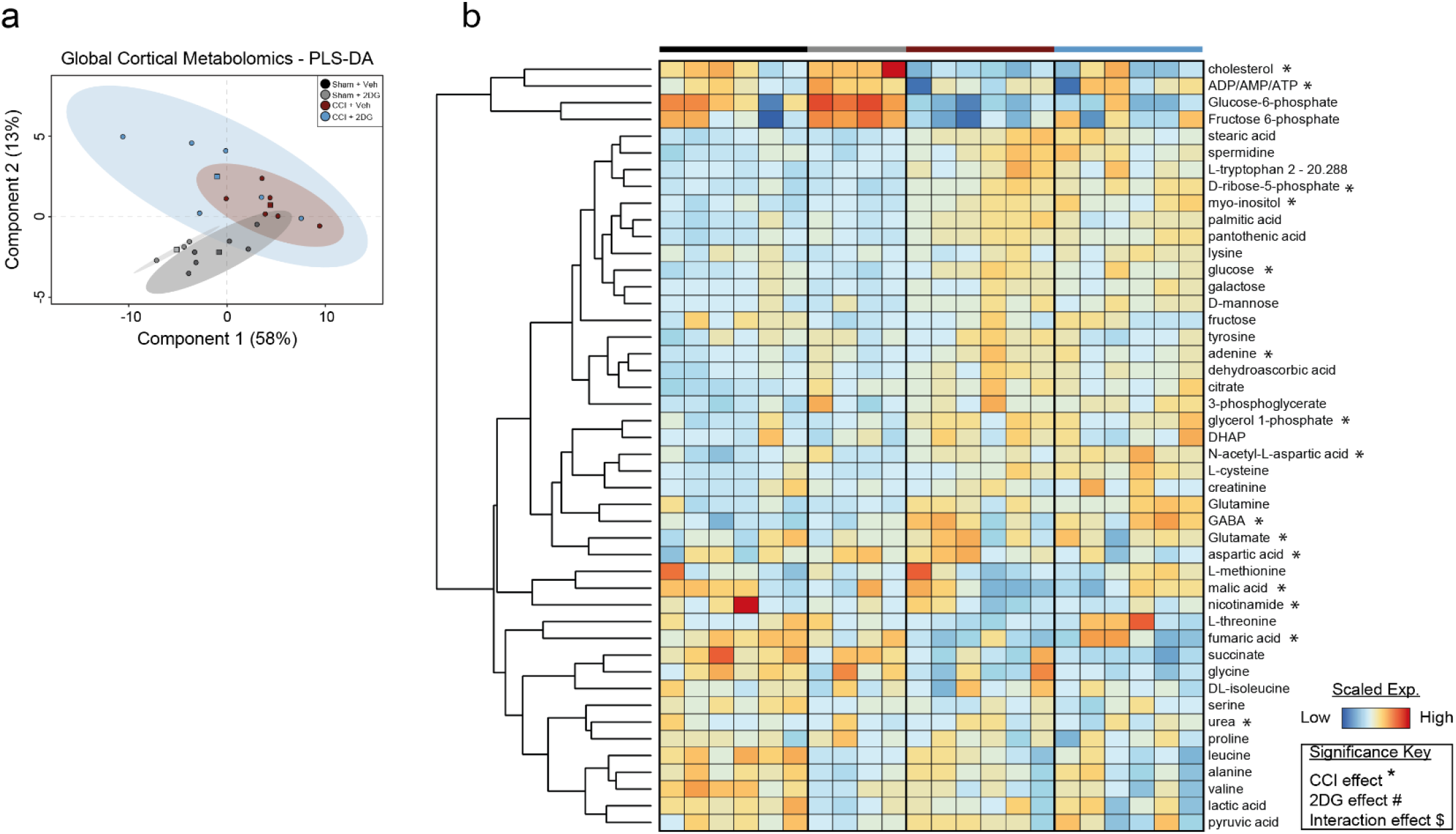
GC-MS analysis of injured and 2DG-treated cortical tissue. **a.** PLS-DA score plot for comparison of the global metabolite profiles of 21 cortical tissue samples - 6 samples from Sham+Veh, 3 samples from Sham+2DG, 6 samples from CCI+Veh, and 6 samples from CCI+2DG. Circles represent individual samples. Squares represent group centroids. Shaded ellipses represent the 95% confidence intervals. **b.** Heatmap illustration of the normalized, scaled expression of each metabolite in each sample. * p < 0.05 by two-way ANOVA with Bonferroni correction.

**Supplementary Figure 4.**
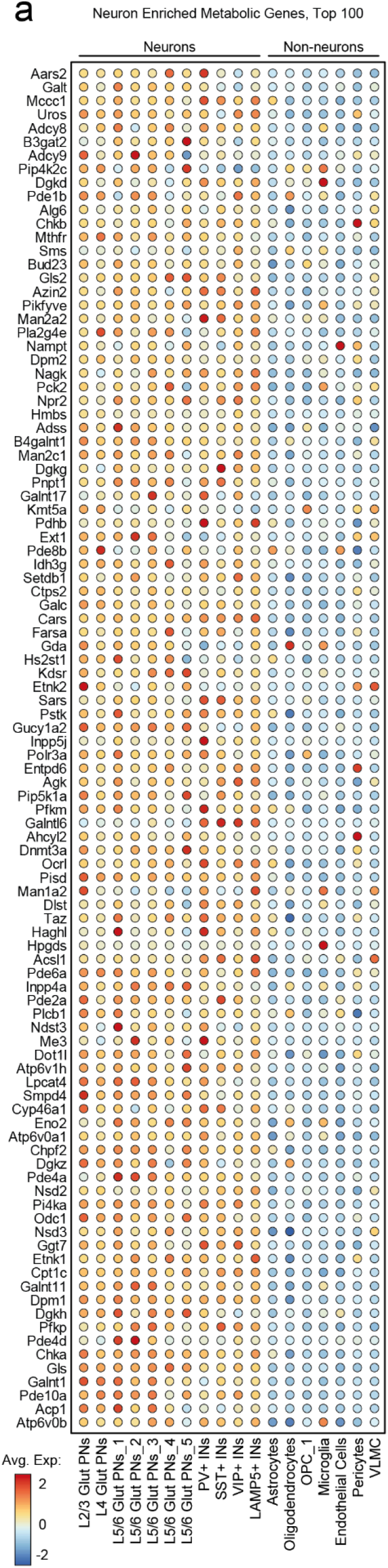
Top neuron and glia enriched metabolic genes. **a.** Dotplot of the top 100 neuron-enriched metabolic genes (inclusion criteria: absolute log_2_FC > 0, p < 0.05 by Wilcoxon rank sum with Bonferroni correction).

**Supplementary Figure 5.**
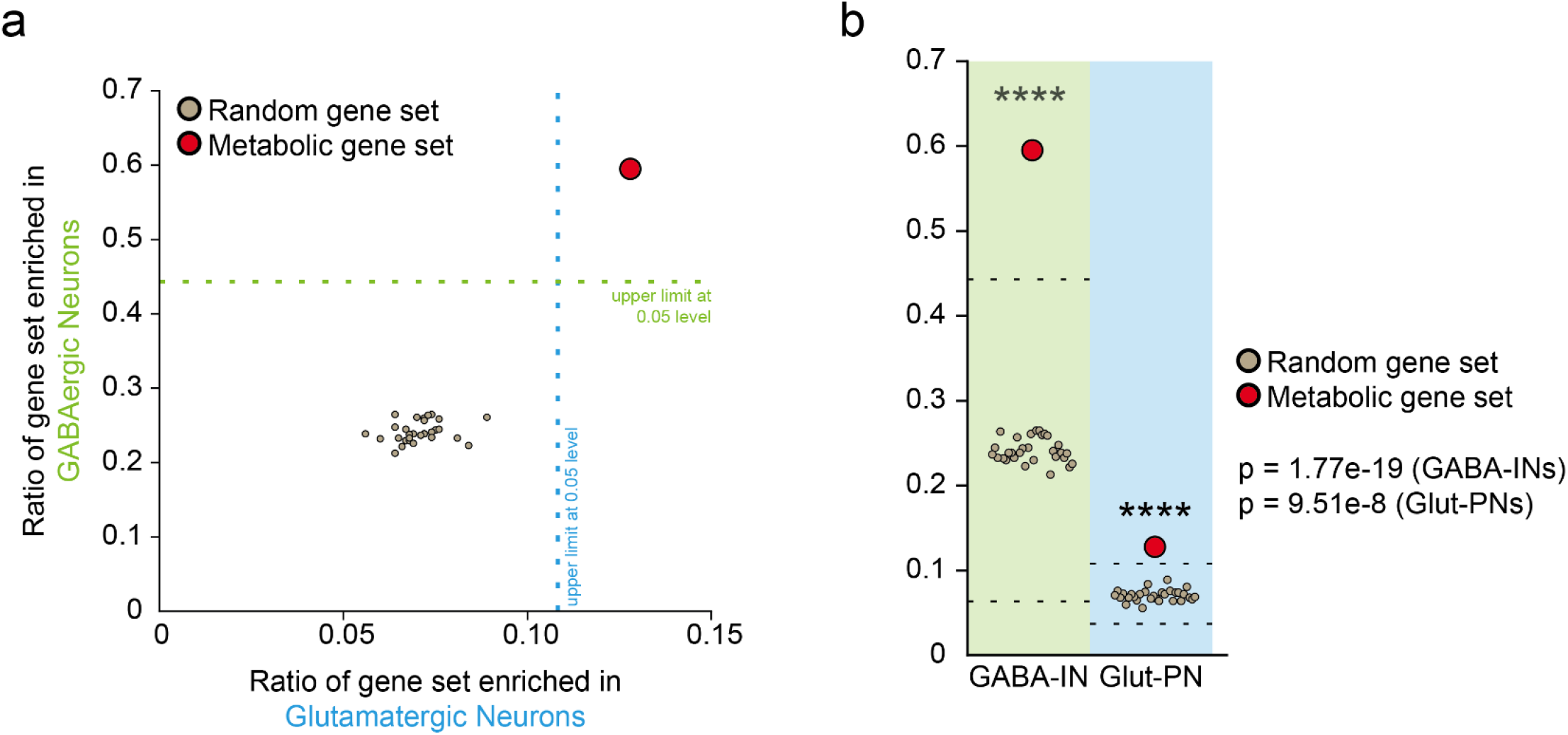
Testing metabolic gene set enrichment profile against random-sampling enrichment profiles. **a.** Scatter plot depiction of the ratio of each gene enriched in Glut-PNs (x-axis) and ratio of each gene enriched in GABA-INs (y-axis) in various gene sets. Each grey dot represents an individual gene set comprised of 1000 random, non-repeating genes. The red dot is the neuron-enriched metabolic gene set from Fig. 4. **b.** Quantification of (**a**.) showing the metabolic gene set is a statistical outlier. The population of random gene sets was normally distributed as determined by Shapiro-Wilks test (GABA-IN enrichment, p = 0.11; Glut-PN enrichment, p = 0.40). **** p < 0.00001 by Grubbs test. Dotted lines represent upper and lower limits at p = 0.05 level.

**Supplementary Figure 6.**
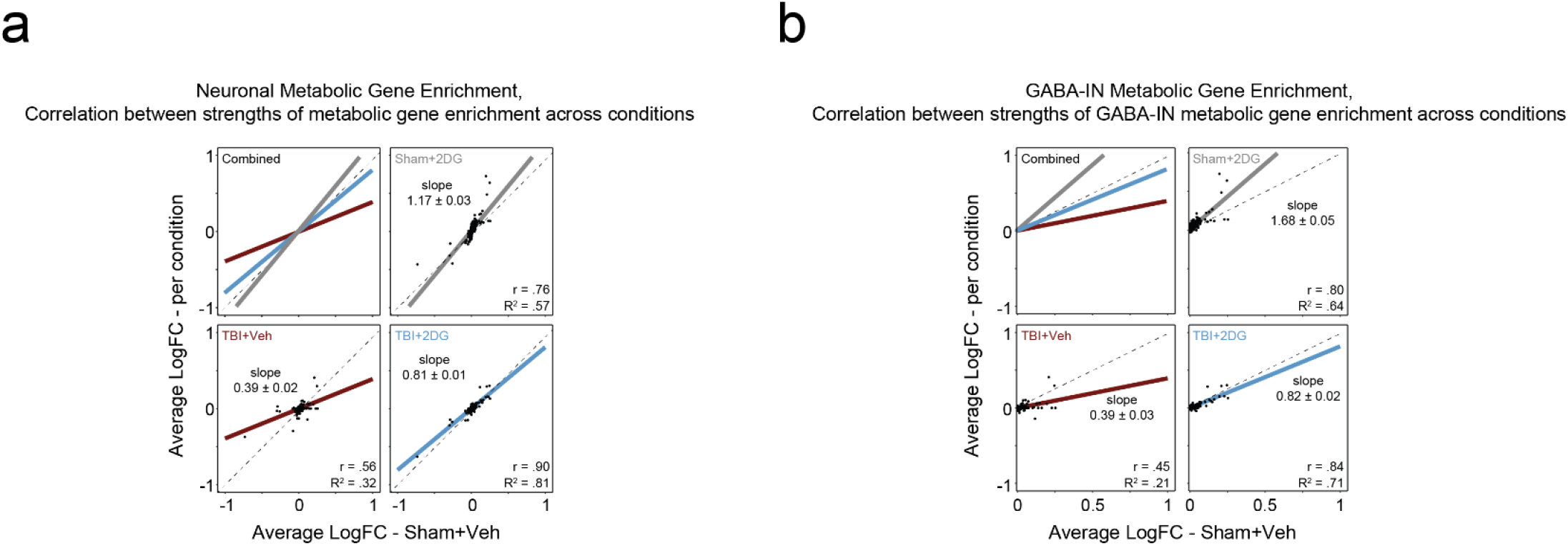
Metabolic gene enrichment correlation plots. **a.** Scatter plots showing correlation between Sham+Veh enrichment signature (x-axis) and each other condition (y-axis). Each dot represents a single neuron-enriched metabolic gene from Fig 4. Dotted line represents a correlation of 1. Colored line represents the linear fit applied to each condition. Negative values represent enrichment in Glut-PNs, positive values represent enrichment in GABA-INs. **b.** Same as (**a**.), except the input genes are only the GABA-IN enriched subset of neuron-enriched metabolic genes.

**Supplementary Figure 7.**
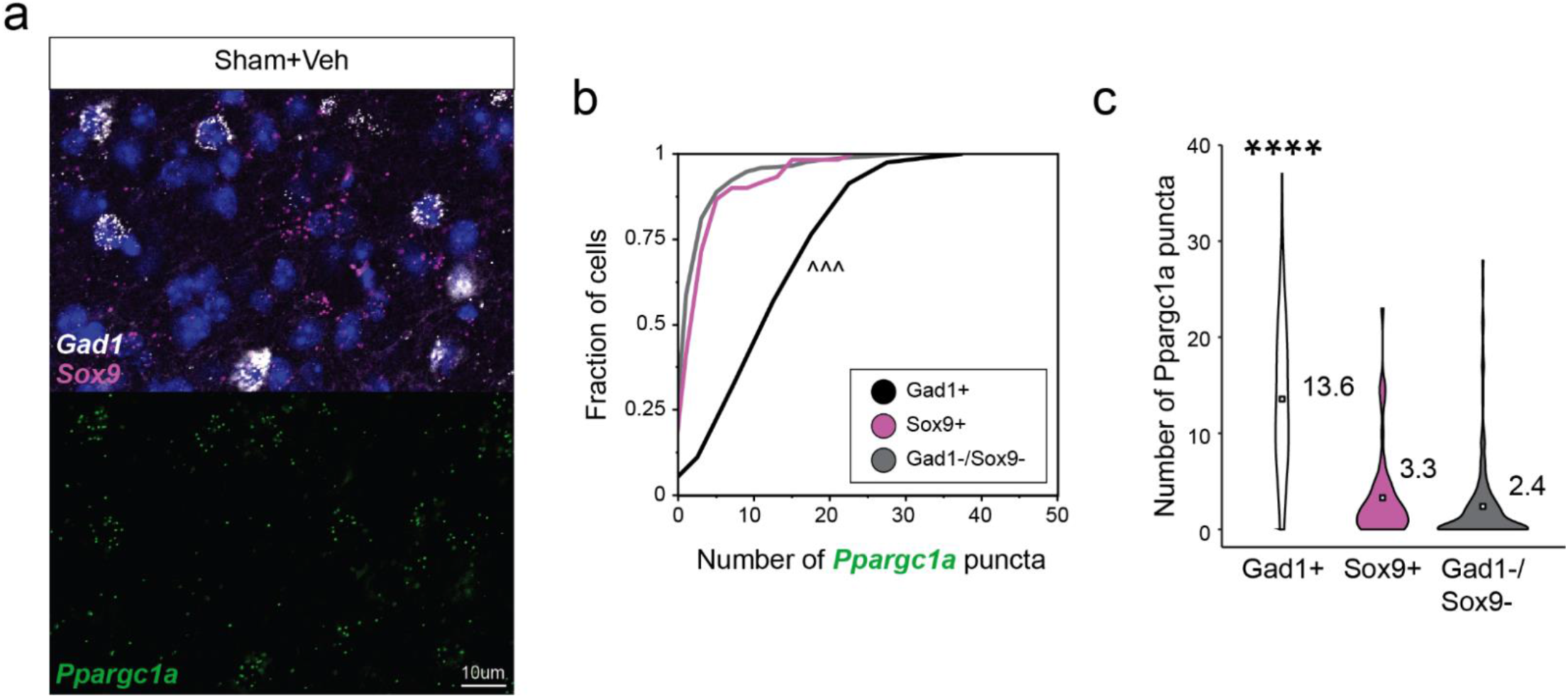
*Ppargc1a* is preferentially expressed and enriched in GABAergic INs. **a**. Representative smFISH image from Sham+Veh in layer 5/6 somatosensory cortex. *Gad1* (white), *Sox9* (magenta), and *Ppargc1a* (green). Scale bar = 10µm. **b.** Cumulative distribution of *Ppargc1a* mRNA puncta count per cell.^^^P < 1 × 10-5 by 2-sample Kolmogorov-Smirnov test with correction for multiple comparison. **c.** Violin plot of *Ppargc1a* mRNA puncta count per cell, inner squares represent mean values. **** p < 0.00001 by one-sided Mann Whitney test with correction for multiple comparison.

**Supplementary Figure 8.**
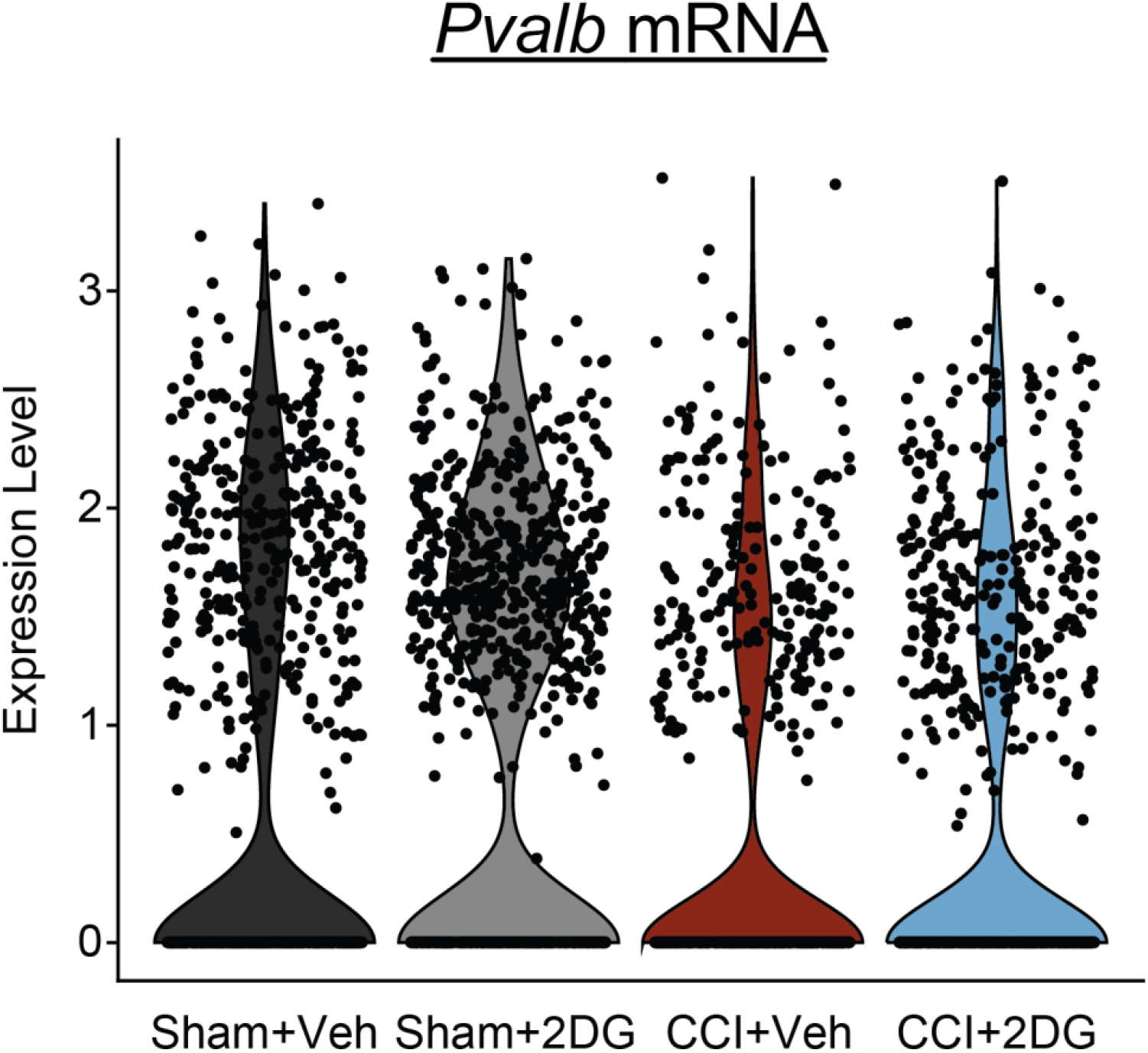
Pvalb mRNA expression is insufficient to stratify PV-INs after injury. Violin plot of *Pvalb* expression in PV-INs across each experimental condition.

**Supplementary Figure 9.**
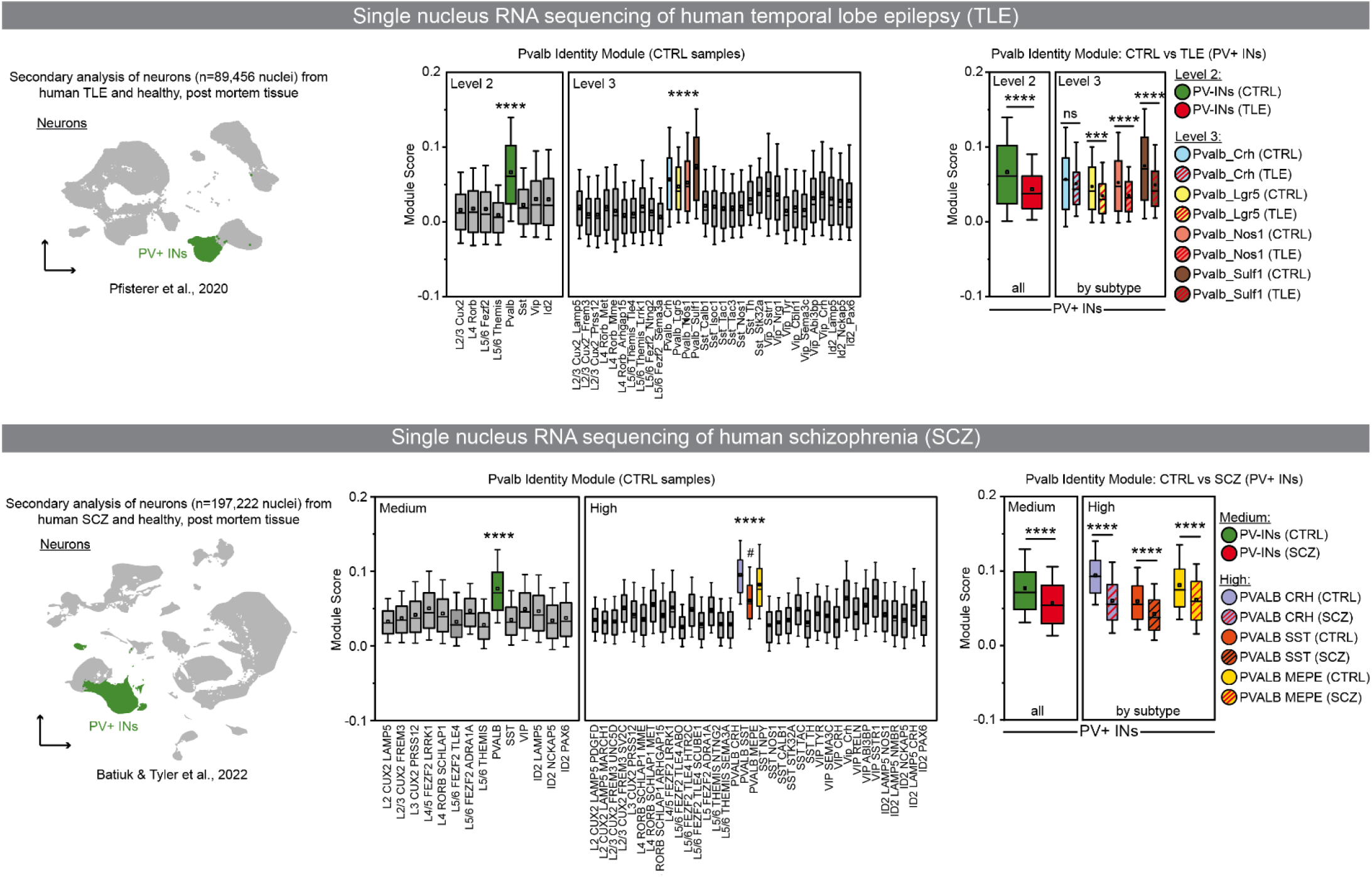
Re-analysis of neurons from published human temporal lobe epilepsy (TLE) and human schizophrenia (SCZ) datasets. **a.** UMAP depiction of reanalyzed neuronal nuclei from published dataset^59^ using human temporal lobe epilepsy (TLE) and post-mortem control (CTRL) (no epilepsy) temporal lobe cortical tissue. PV-INs are highlighted in green. **b.** Boxplot of Pvalb identity module scores across neurons in the CTRL samples showing significant enrichment of Pvalb Identity in PV-INs compared to all other neuron groups. The left panel groups neurons into a lower tier of classification, ‘level 2’, while the right panel groups neurons into a higher tier of classification, ‘level 3’, in which there are 4 PV-IN subtypes. **** p < 0.0001 by Wilcoxon Rank Sum with Bonferroni correction **c.** Boxplot of Pvalb identity module scores across PV-INs in CTRL and TLE samples, level 2 (left) and level 3 (right). *** p < 0.001, **** p < 0.0001 by Wilcoxon Rank Sum with Bonferroni correction. **d.** UMAP depiction of reanalyzed neuronal nuclei from published dataset^60^ using human schizophrenia (SCZ) and post-mortem CTRL (no schizophrenia) dorsal-lateral prefrontal cortical tissue. PV-INs are highlighted in green. **e.** Boxplot of Pvalb identity module scores across neurons in the CTRL samples showing significant enrichment of Pvalb Identity in PV-INs compared to all other neuron groups. The left panel groups neurons into a lower tier of classification, ‘medium’, while the right panel groups neurons into a higher tier of classification, ‘high’, in which there are 3 PV-IN subtypes. **** p < 0.0001 by Wilcoxon Rank Sum with Bonferroni correction. All PV-IN subtypes scored significantly higher (p<0.0001) than non-PVALB neurons except for Pvalb_SST, denoted by #, that did not score higher than VIP_Crh or VIP_SSTR1. **f.** Boxplot of Pvalb identity module scores across PV-INs in CTRL and SCZ samples, medium (left) and high (right). *** p < 0.001, **** p < 0.0001 by Wilcoxon Rank Sum with Bonferroni correction.

